# The lncRNA *APOLO* interacts with the transcription factor WRKY42 to trigger root hair cell expansion in response to cold

**DOI:** 10.1101/2020.07.13.188763

**Authors:** Michaël Moison, Javier Martínez Pacheco, Leandro Lucero, Camille Fonouni-Farde, Johan Rodríguez-Melo, Natanael Mansilla, Aurélie Christ, Jérémie Bazin, Moussa Benhamed, Fernando Ibañez, Martin Crespi, José M. Estevez, Federico Ariel

**Affiliations:** Instituto de Agrobiotecnología del Litoral, Universidad Nacional del Litoral, CONICET, FBCB/FHUC, Centro Científico Tecnológico CONICET Santa Fe, Colectora Ruta Nacional No 168 km. 0, Paraje El Pozo, Santa Fe 3000, Argentina; Fundación Instituto Leloir and IIBBA-CONICET, Av. Patricias Argentinas 435, Buenos Aires CP C1405BWE, Argentina; Instituto de Investigaciones Agrobiotecnológicas, CONICET, Universidad Nacional de Río Cuarto, Río Cuarto 5800, Argentina; Institute of Plant Sciences Paris-Saclay (IPS2), CNRS, INRA, University Paris-Saclay and University of Paris Bâtiment 630, 91192 Gif sur Yvette, France; Centro de Biotecnología Vegetal (CBV), Facultad de Ciencias de la Vida (FCsV), Universidad Andres Bello, Santiago, Chile and Millennium Institute for Integrative Biology (iBio), Santiago, Chile

**Author notes:** Correspondence to: Federico Ariel and José M. Estevez. These authors contributed equally to this work (Michaël Moison, Javier Martínez Pacheco, Leandro Lucero).

**Keywords:** root hairs, long noncoding RNAs, *APOLO*, RHD6, WRKY42, cold temperature

## Abstract

Plant long noncoding RNAs (lncRNAs) have emerged as important regulators of chromatin dynamics, impacting on transcriptional programs leading to different developmental outputs. The lncRNA *AUXIN REGULATED PROMOTER LOOP* (*APOLO*) directly recognizes multiple independent loci across the *Arabidopsis* genome and modulates their three-dimensional chromatin conformation, leading to transcriptional shifts. Here, we show that *APOLO* recognizes the locus encoding the root hair (RH) master regulator ROOT HAIR DEFECTIVE 6 (RHD6) and controls *RHD6* transcriptional activity leading to cold-enhanced RH elongation through the consequent activation of the transcription factor gene RHD6-like *RSL4*. Furthermore, we demonstrate that *APOLO* interacts with the transcription factor WRKY42 and modulates its binding to the *RHD6* promoter. WRKY42 is required for the activation of *RHD6* by low temperatures and *WRKY42* deregulation impairs cold-induced RH expansion. Collectively, our results indicate that a novel ribonucleoprotein complex involving *APOLO* and WRKY42 forms a regulatory hub which activates *RHD6* by shaping its epigenetic environment and integrates signals governing RH growth and development.

**SUMMARY:** The lncRNA *APOLO* directly regulates the transcription of the root hair-master gene *RHD6*. In response to cold, *APOLO* is induced and it decoys the H3K27me3-binding protein LHP1 away from *RHD6*. In addition, *APOLO* modulates the binding of the transcription factor WRKY42 to the *RHD6* promoter at low temperatures.

## INTRODUCTION

Root hairs (RHs) are single cell projections developed from specialized epidermal trichoblast cells able to increase their size several hundred times in a polar manner to reach and promote the uptake of water-soluble nutrients, interact with soil microorganisms and support anchor to the plant. The specification of epidermal cells into RHs is a complex process whose underlying mechanisms are partially understood. In *Arabidopsis thaliana*, RH cell fate is controlled by a developmental program involving a complex of transcription factors (TFs) promoting the expression of the homeodomain protein GLABRA 2 (GL2) (Ryu et al., 2005; Song et al., 2011; Schiefelbein et al., 2014; Balcerowicz et al., 2015). GL2 blocks RH development by inhibiting the transcription of the master regulator *ROOT HAIR DEFECTIVE 6* (*RHD6*) (Lin et al., 2015). In the cells that differentiate into RHs (known as trichoblasts), a second TF complex suppresses *GL2* expression (Schiefelbein et al., 2014), forcing the cells to enter the RH cell fate program via the concomitant activation of RHD6 along with downstream TFs (Menand et al., 2007; Pires et al., 2013). Briefly, RHD6 together with its homolog RSL1 (ROOT HAIR DEFECTIVE 6 LIKE 1) induce the expression of TFs from the bHLH family, including RSL2 (ROOT HAIR DEFECTIVE 6 LIKE 2) and RSL4 (ROOT HAIR DEFECTIVE 6 LIKE 4), ultimately triggering the differentiation of the RHs and their subsequent polarized tip-growth (Karas et al., 2009; Yi et al., 2010; Bruex et al., 2012). In addition, it was proposed that RSL4 controls the expression of a small subset of nearly 125 genes (Won et al., 2009; Yi et al., 2010; Datta et al., 2015; Vijayakumar et al., 2016), including several cell wall extensins (EXTs) (Ringli, 2010; Velasquez et al., 2011) sufficient to promote RH growth (Hwang et al., 2017).

RH expansion is regulated both by cell-intrinsic factors (e.g. endogenous phytohormones such as auxin) and external environmental signals (e.g. phosphate (Pi) availability in the soil) (Mangano et al., 2017; Bhosale et al., 2018). Pi starvation is one of the key environmental factors promoting rapid RH growth (Yi et al., 2010; Datta et al., 2015; Vijayakumar et al., 2016). In *Arabidopsis*, it triggers *RSL4* expression via an enhanced auxin production, activating downstream effector genes mediating cell growth (Yi et al., 2010; Datta et al., 2015; Mangano et al., 2017; Marzol et al., 2017; Bhosale et al., 2018). Accordingly, several auxin-related TFs have been implicated in Pi-starvation signaling in roots, including WRKY proteins that control the expression of the Pi transporter families Pi-permease PHO1 and PHOSPHATE TRANSPORTER (PHT) (Devaiah et al., 2007; Chen et al., 2009; Wang et al., 2014; Su et al., 2015). Under Pi-sufficient conditions, WRKY6 and WRKY42 bind to W-boxes of the *PHO1* promoter and suppress its expression. During Pi starvation, WRKY42 is degraded by the 26S proteasome pathway, resulting in the activation of *PHO1* transcription (Chen et al., 2009; Su et al., 2015). In addition, WRKY42 functions as a positive regulator of *PHT1;1*, by binding to its promoter under Pi-sufficient condition (Su et al., 2015). Overall, WRKY42 is part of the components activating root early-responses to Pi starvation, although its role in controlling RH growth remains unexplored.

In recent years, plant long noncoding RNAs (lncRNAs) have emerged as important regulators of gene expression, and several among them, have been functionally linked to Pi homeostasis. For instance, the lncRNA *INDUCED BY PHOSPHATE STARVATION 1* (*IPS1*) can sequester the Pi starvation-induced microRNA miR-399, attenuating miR-399-mediated repression of *PHO2*, a gene encoding an E3 ligase affecting Pi uptake (Franco-Zorrilla et al., 2007). In addition, the *cis*-natural antisense (*cis*-NAT) transcript *PHO1;2*, induced under Pi deficiency, was shown to promote the translation of the *PHO1;2* mRNA involved in Pi loading into the xylem. The expression of this *cis*-NAT is associated with the transport of the sense–antisense RNA pair toward the polysomes (Jabnoune et al., 2013). More recently, it was shown that the lncRNA *AUXIN REGULATED PROMOTER LOOP* (*APOLO*) recognizes multiple spatially independent genes by sequence complementarity and DNA-RNA duplex formation, known as R-loops. Upon recognition, *APOLO* shapes the three-dimensional (3D) conformation of its target regions by decoying the Polycomb Repressive Complex 1 (PRC1) component LIKE HETEROCHROMATIN PROTEIN 1 (LHP1), thereby regulating their transcription (Ariel et al., 2014; Ariel et al., 2020).

Here, we show that the lncRNA *APOLO* directly regulates a subset of genes involved in RH development, including the master regulator of RH initiation *RHD6*. *APOLO* activates *RHD6* transcription by modulating the formation of a local chromatin loop encompassing its promoter region, an epigenetic regulatory mechanism likely involving PRC1 and PRC2 components. Furthermore, we found that *APOLO* interacts with the TF WRKY42, forming a new hub that regulates *RHD6* to induce RH growth in response to low temperatures. RHD6-mediated induction of RH expansion likely occurs through the transcriptional activation of the TF-encoding gene *RSL4*, which emerged as a key factor in the response to cold.

## RESULTS

### *APOLO* regulates root hair cell elongation in response to low temperatures

Based on Chromatin Isolation by RNA Purification (ChIRP-Seq) performed in wild-type (WT, Col-0) plants, it was previously reported that the lncRNA *APOLO* recognizes a subset of independent loci enriched in categories related to cell wall composition and organization (Ariel et al., 2020). A closer look at *APOLO bona fide* targets allowed us to identify seventeen genes involved in RH growth and expansion (**Supplementary Table 1**), a process dependent on cell wall remodeling molecules, including EXTs and EXT-related proteins (Ringli, 2010; Lamport et al., 2011; Velasquez et al., 2011; Velasquez et al., 2015; Marzol et al., 2018). Interestingly, according to single-cell RNA-seq datasets (Zhang et al., 2019) *APOLO* transcripts are enriched in RH cells (**Supplementary Figure 1A**). Notably, sixteen *APOLO* direct targets were upregulated upon *APOLO* over-expression (**Supplementary Table 2**). Furthermore, 52 additional RH-related genes were upregulated in *35S:APOLO* seedlings (**Supplementary Table 2**) (Ariel et al., 2020). Among them, the RH central TFs *RHD6* (as a direct target), *RSL2* and *RSL4* (as indirectly regulated) were induced upon *APOLO* over-expression.

It was reported that the *APOLO* locus is targeted by the RNA-polymerase Pol V and silenced by RNA-directed DNA Methylation (RdDM,(Ariel et al., 2014)). A search in a small RNA-Seq performed in WT roots subjected to different temperature treatments (Gyula et al., 2018) revealed that RdDM-related 24nt siRNA accumulation over the *APOLO* locus is less abundant at low temperatures (**Supplementary Figure 1B**), suggesting that *APOLO* transcription is regulated by cold. Accordingly, we found that *APOLO* transcriptional accumulation increases in roots after 24h at 10°C (**Figure 1A**). An analysis of the promoter activity of the 5.2kb region upstream *APOLO* (Ariel et al., 2020) directing the expression of a *GFP* reporter gene, additionally revealed a higher transcriptional activity at low temperatures in the RHs (**Figure 1B**). Strikingly, we observed that two RNAi-*APOLO* (repression of approximately 90%; Ariel et al., 2014) and *35S:APOLO* independent lines (over expression of 30-fold and 60-fold, respectively; Ariel et al., 2020) exhibit a basal increase of RH length at 22°C, and uncovered a strong induction of RH elongation in WT and RNAi-*APOLO* at 10°C, in contrast to *35S:APOLO* lines (**Figure 1C**). Accordingly, *RHD6* is induced in response to cold in WT roots, whereas RNAi-*APOLO* and *35S:APOLO* roots display higher *RHD6* basal levels than the WT (**Figure 1D**). Notably, *RHD6* transcript levels are not further induced by cold in *35S:APOLO* roots (**Figure 1D**), in agreement with the RH phenotype (**Figure 1C**). Collectively, our findings suggest that *APOLO* participates in the induction of cold-mediated RH elongation and a deregulation of *APOLO* transcript levels can impact RH growth.

**Figure 1.**
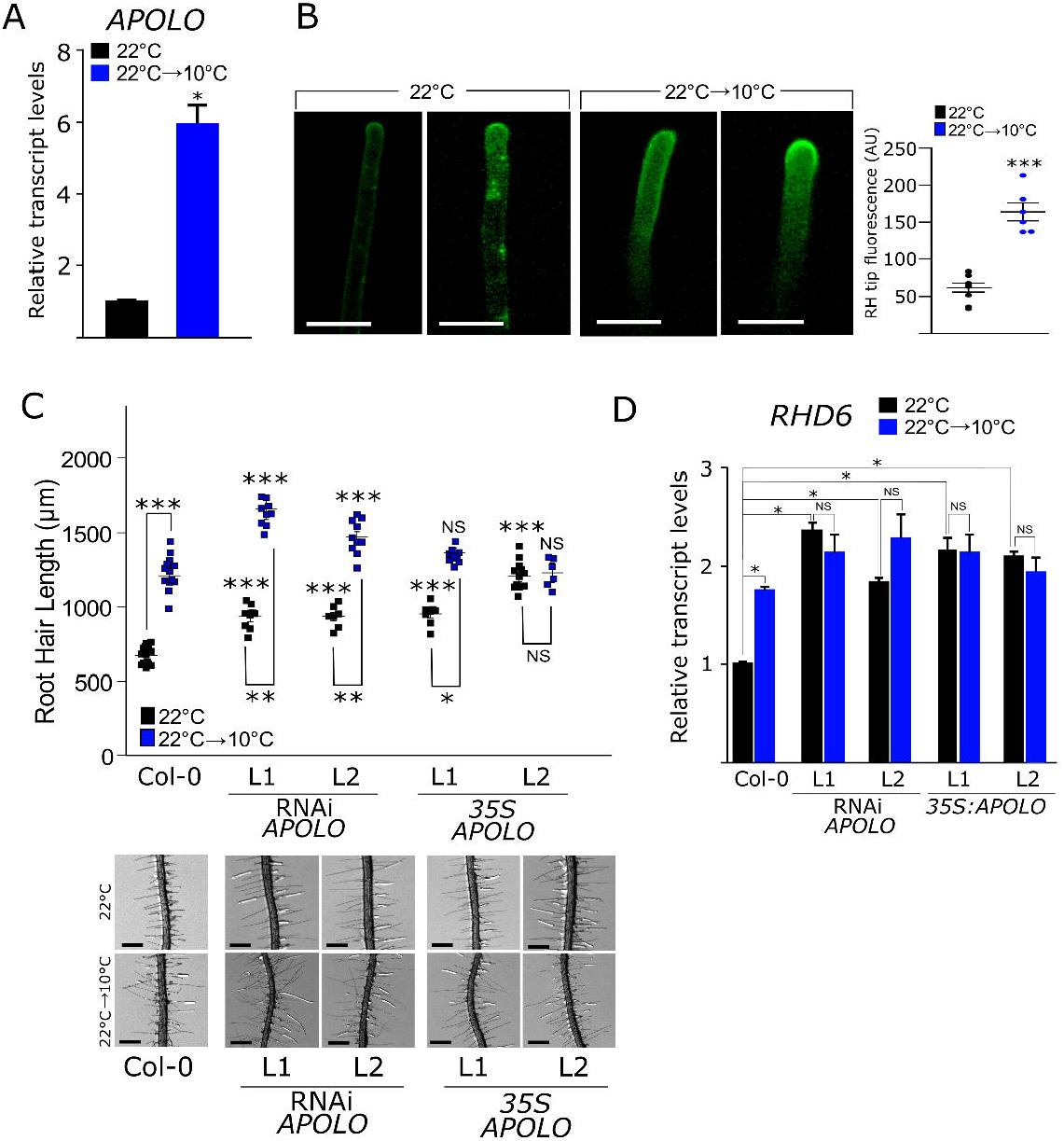
*APOLO* regulates root hair elongation in response to low temperatures. **A.** *APOLO* transcript levels measured by RT-qPCR in roots of 10-day-old plants grown at 22 °C treated or not for 24h at 10 °C. **B.** The *APOLO* promoter region (full intergenic region between *PID* and *APOLO*) is active in root hairs at 22 °C and 10 °C, using the *GFP* reporter gene. Images are representative of the lowest and highest GFP signal detected in the observation of several roots. The quantification of the fluorescence is shown in the right panel. **C.** Quantification of RH length of 8-day-old Col-0, RNAi-*APOLO* and *35S:APOLO* plants, respectively, at 22 °C and 10 °C. Each point is the mean ±error of the length of the 10 longest RHs identified in the root elongation zone in a single root. Representative images of each genotype are shown below the graph. The scale bas represent 500μm. Asterisks indicate significant differences between Col-0 and the corresponding genotype at the same temperature or significant differences between temperature treatments on the same genotype at the same temperature (one way ANOVA followed by a Tukey-Kramer test; “*”<0.05, “**”<0.01, “***”<0.001). NS stands for no statistically significant difference. **D.** *RHD6* transcript levels measured by RT-qPCR in roots of Col-0 plants vs. two independent RNAi-*APOLO* lines and two independent *35S:APOLO* lines grown at 22 °C treated or not for 24h at 10 °C. In A and D, the error bars represent the SD of 3 biological replicates. The asterisks indicate that the differences are significant (t test p<0.05). Values are normalized using the constant housekeeping transcript *PP2A*.

Previous studies pointed out a key role of RHD6 (together with RSL1) in RH development, which is mediated by RSL4 and RSL2 as downstream regulators of RH cell elongation (Menand et al., 2007; Pires et al., 2013). Notably, *RHD6*, *RSL2* and *RSL4* transcript levels were upregulated in *35S:APOLO* seedlings (**Supplementary Table 2**; (Ariel et al., 2020)), although only *RHD6* was identified as a direct target of *APOLO* (Ariel et al, 2020). In agreement with *RHD6* transcriptional behavior (**Figure 1D**), *RSL2* and *RSL4* basal transcript levels are higher in RNAi-*APOLO* and *35S:APOLO* compared to WT. Interestingly, *RSL2* and *RSL4* are induced by cold in WT and RNAi-*APOLO*, but not in *35S:APOLO* roots (**Supplementary Figure 2A**), suggesting that low temperatures can activate these two genes still in the absence of *APOLO* and bypassing RHD6 for RH expansion (**Figure 1C**). Thus, we assessed if these TFs were also controlling the promotion of RH growth by low temperatures. To this end, we tested how *rhd6/rsl1/rsl4* and *rsl2*, *rsl4* and double mutant plants *rsl2/rsl4* respond to low temperatures in comparison with control conditions **(Supplementary Figure 2A)**. The *rsl2* mutant was highly responsive to low temperatures in a similar manner to WT while *rsl4* was impaired in the response to cold. The double mutant *rsl2/rsl4* and the triple mutant *rhd6/rsl1/rsl4* did not develop RHs in either of the two conditions. In addition, constitutive expression of *RSL4* (*35S:RSL4*) as well as its expression under the control of the RH specific *EXPANSIN7* promoter (*EXP7p:RSL4*) boosted basal RH growth without further enhancement in response to cold (**Supplementary Figure 2B**). These results demonstrate that RSL4 is a key factor mediating RHD6 activation of RH growth at low temperature, and RSL2 might participate to a lower extent.

Nutrient unavailability is known to activate RH expansion through a transcriptional reprogramming governed by RHD6 and downstream TFs. The quantification of RH growth of WT plants in response to increasing concentrations of nutrients (0.5X to 2.0X MS (Murashige and Skoog) medium) indicates that high concentrations impair RH growth triggered by low temperatures **(Supplementary Figure 3A**). In a similar way, an increase in agar concentration in the MS medium (from 0.8% to 2.5%), which likely re-strains nutrient mobility (Singha, S., Townsend, E.C. and Oberly, 1985; Nonami and Boyer, 1989; Ghashghaie et al., 1991; Buah et al., 1999) blocks cold-induced RH expansion **(Supplementary Figure 3B**). Altogether, these observations suggest that low temperatures restrict nutrient mobility and availability in the culture medium, leading to the promotion of polar RH growth.

### *APOLO* directly modulates the three-dimensional chromatin conformation of the root hair specific locus *RHD6*

Among *APOLO* targets involved in RH development, we found the master regulator of RH initiation *RHD6* (Menand et al., 2007; Pires et al., 2013). The epigenetic profile of the *RHD6* locus corresponds to typical *APOLO* targets (**Figure 2A**, (Ariel et al., 2020)), including H3K27me3 deposition (track 1), LHP1 recognition (track 2, chromatin immunoprecipitation (ChIP)-Seq, (Veluchamy et al., 2016)), and *APOLO* binding regions (tracks 3 to 5, chromatin isolation by RNA purification (ChIRP)-Seq, (Ariel et al., 2020)). A GAAGAA box, shown to be important for *APOLO* target recognition (Ariel et al., 2020) is located in the *RHD6* locus and coincides with *APOLO* binding site. In addition, a peak of DNA-RNA hybrid immunoprecipitation (DRIP)-Seq from root samples indicates the presence of an R-loop coinciding with *APOLO* recognition sites over *RHD6* (tracks 6 to 8, (Xu et al., 2020)).

**Figure 2.**
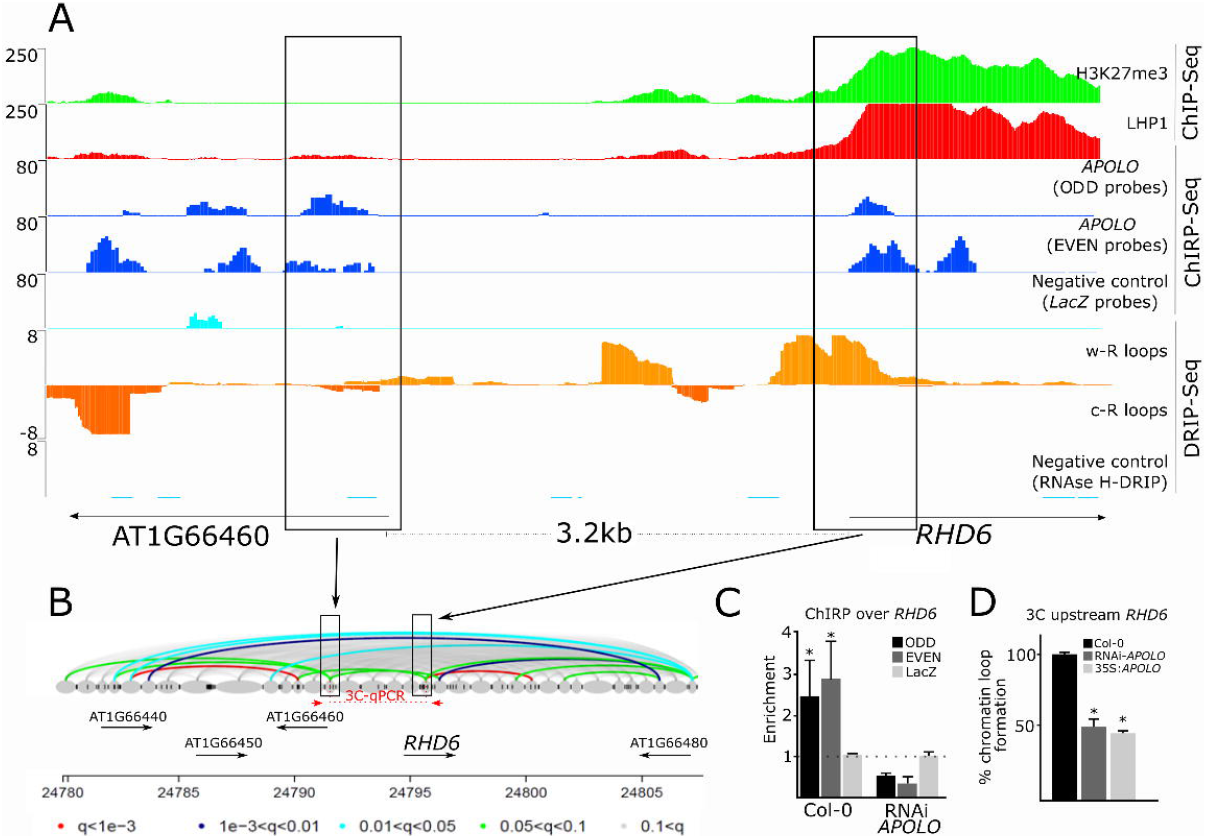
*APOLO* directly modulates chromatin three-dimensional conformation of the *RHD6* locus. **A.** Epigenomic landscape of the *RHD6* locus. Lane 1: H3K27me3 deposition by ChIP-Seq (Veluchamy et al., 2016). Lane 2: LHP1 deposition by ChIP-Seq (Veluchamy et al., 2016). Lane 3 to 5: *APOLO* recognition by ChIRP-Seq (Lane 3 and 4, using ODD and EVEN sets of probes against *APOLO*, respectively; Lane 5, negative control using LacZ probes)(Ariel et al., 2020). Lane 6 to 8: R-loop formation by DRIP-Seq (R-loop Atlas, root samples,(Xu et al., 2020), on Watson strand (Lane 6) and Crick strand (Lane 7). DRIP negative control after RNAseH treatment is shown in Lane 8. Gene annotation is shown at the bottom. **B.** Chromatin loops identified in the *RHD6* region by *Dpn*II HiC (Liu et al., 2016). Colors of the loops are related to the corresponding q-values indicated below. Black boxes in A and B indicate the same genomic locations, where the bases of the chromatin loop correlate with R-loop formation and *APOLO* recognition (compared to gene annotations). In red the arrows indicate the position of the probes used for 3C-qPCR shown in **D**. **C.** *APOLO* association to DNA of the *RHD6* locus by ChIRP-qPCR in Col-0 and RNAi-*APOLO* plants. The background level was determined using a set of probes against LacZ RNA. **D.** Relative chromatin loop formation by 3C-qPCR deduced from **B**, in Col-0 plants vs. *35S:APOLO* and RNAi lines for the region upstream *RHD6*. The probes used for 3C-qPCR are indicated in red in panel **B**. In C and D, the error bars represent the SD of 3 biological replicates. The asterisks indicate that the differences are significant (t test “*”<0.05, “***”<0.001).

Remarkably, *APOLO* recognition and R-loop formation are also detectable over *RHD6* neighbor gene, located 3.2 kb upstream *RHD6* transcription start site (**Figure 2A**). According to *Dpn*II Hi-C datasets from Arabidopsis seedlings (Liu et al., 2016), a chromatin loop encompassing the intergenic region upstream *RHD6* was detected (**Figure 2B**), and coincides with *APOLO* binding-sites (**Figure 2A**, ChIRP-Seq). By performing a ChIRP-qPCR with two independent sets of biotinylated probes to purify *APOLO* (ODD and EVEN; (Ariel et al., 2020)) and one additional set used as a negative control (LacZ), we confirmed that *APOLO* RNA–*RHD6* DNA interaction occurs in WT and is lost in *APOLO* knockdown (RNAi) seedlings ((Ariel et al., 2014); **Figure 2C**). In addition, the quantification of relative *RHD6* loop formation in RNAi-*APOLO* and *35S:APOLO* (Ariel et al., 2020) seedlings, revealed impaired loop formation in both lines (**Figure 2D**), hinting at a stoichiometric requirement of *APOLO* for *RHD6* chromatin loop formation. Chromatin loop formation (**Figure 2D**) is in agreement with *RHD6* basal levels in *35S:APOLO* and RNAi-*APOLO* lines (**Figure 1D**), suggesting that the chromatin loop including *RHD6* promoter region precludes transcription. Altogether, our results indicate that *APOLO* lncRNA directly regulates *RHD6* transcriptional activity by fine-tuning local chromatin 3D conformation.

It was previously reported that PRC2 actively participates in the regulation of RH growth (Ikeuchi et al., 2015) and that the *RHD6* locus exhibits H3K27me3 deposition and LHP1 recognition (**Figure 2A;** (Veluchamy et al., 2016)). Considering that the lncRNA *APOLO* interacts with the PRC1 component LHP1 *in vivo* (Ariel et al., 2014; Ariel et al., 2020), we decided to explore the role of PRC1 and PRC2 in *APOLO*-mediated *RHD6* activation at low temperatures. At 22°C, *RHD6* suffers a reduction of H3K27me3 in the PRC2 mutant *curly leaf* (*clf*), in contrast to the PRC1 mutant *lhp1* (**Supplementary Figure 4A;** (Veluchamy et al., 2016)). Interestingly, we observed that H3K27me3 deposition and LHP1 binding diminish in WT roots treated for 24h at 10°C compared to 22°C (**Supplementary Figure 4B**), consistent with the induction of *RHD6* in response to cold (**Figure 2E**). Moreover, *lhp1* and *clf* mutants exhibit a basal decrease of RH length together with a slight decrease of cold-induced RH elongation in *lhp1*, and a strong decrease of cold-induced RH elongation in *clf* (**Supplementary Figure 4D**). Consistently, although the decrease in H3K27me3 deposition results in higher basal transcript levels of *RHD6* in the *clf* background, *RHD6* transcriptional activation by cold is abolished in the *clf* and *lhp1* mutants (**Supplementary Figure 4C**), hinting at an important role of chromatin rearrangement for *RHD6* activation in response to cold.

### *APOLO* interacts with the transcription factor WRKY42 to coordinate the activation of *RHD6*

In order to uncover novel actors involved in cold-induced transcriptional regulation of RH growth, we aimed at identifying *APOLO* protein partners. To this end, we performed an exploratory ChIRP on WT seedlings using two independent set of biotinylated probes to purify *APOLO* (ODD and EVEN; (Ariel et al., 2014; Ariel et al., 2020)), and one additional set used as a negative control (against LacZ), as recently described (Rigo et al., 2020). Co-purified proteins were precipitated and analyzed by mass spectrometry. Among the potential *APOLO* partners (i.e. identified with at least two hits in ODD and EVEN samples, but absent in LacZ-ChIRP), we found the WRKY42 protein, a TF involved in the response to Pi starvation (Su et al., 2015), an environmental condition that promotes RH cell expansion (Bhosale, *et al.*, 2018) in a similar manner to low temperatures. Thus, *APOLO*-WRKY42 interaction was validated by RNA immuno-precipitation (RIP-qPCR) in tobacco leaves and in *Arabidopsis* plants transitory or stably transformed with 35S:*WRKY42:GFP*, respectively (**Figure 3A**). Interestingly, according to the Arabidopsis eFP Browser (Waese et al., 2017), *WRKY42* is induced in roots when seedlings are subjected to 4°C for 24h (**Figure 3B**). At 10°C, we observed that *WRKY42* transcriptional accumulation augments significantly in roots (**Figure 3C**). Notably, 13 out of the 17 *APOLO* targets contained between 1 and 4 canonical WRKY TF binding sites (W-box) in their promoters, including *RHD6* (**Supplementary Table 1**). By using a *35S:WRKY42:GFP* line, we determined that WRKY42 can directly bind to the promoter region of *RHD6* (**Figure 3D**). Accordingly, the over-expression of *WRKY42* (*35S:WRKY42:GFP* line) led to a basal increase of *RHD6* levels (**Figure 3E**) and RH elongation (**Figure 3F**) at ambient temperature, mimicking the effect of cold. On the contrary, cold-mediated induction of *RHD6* is abolished in the *wrky42* mutant ((Su et al., 2015); **Figure 3E**), which consistently exhibits shorter RHs at 22°C and almost no RH elongation at low temperatures (**Figure 3F**). Taken together, these results suggest that the *APOLO*-interacting TF WRKY42 is an important regulator of *RHD6*-mediated RH growth in response to cold.

**Figure 3.**
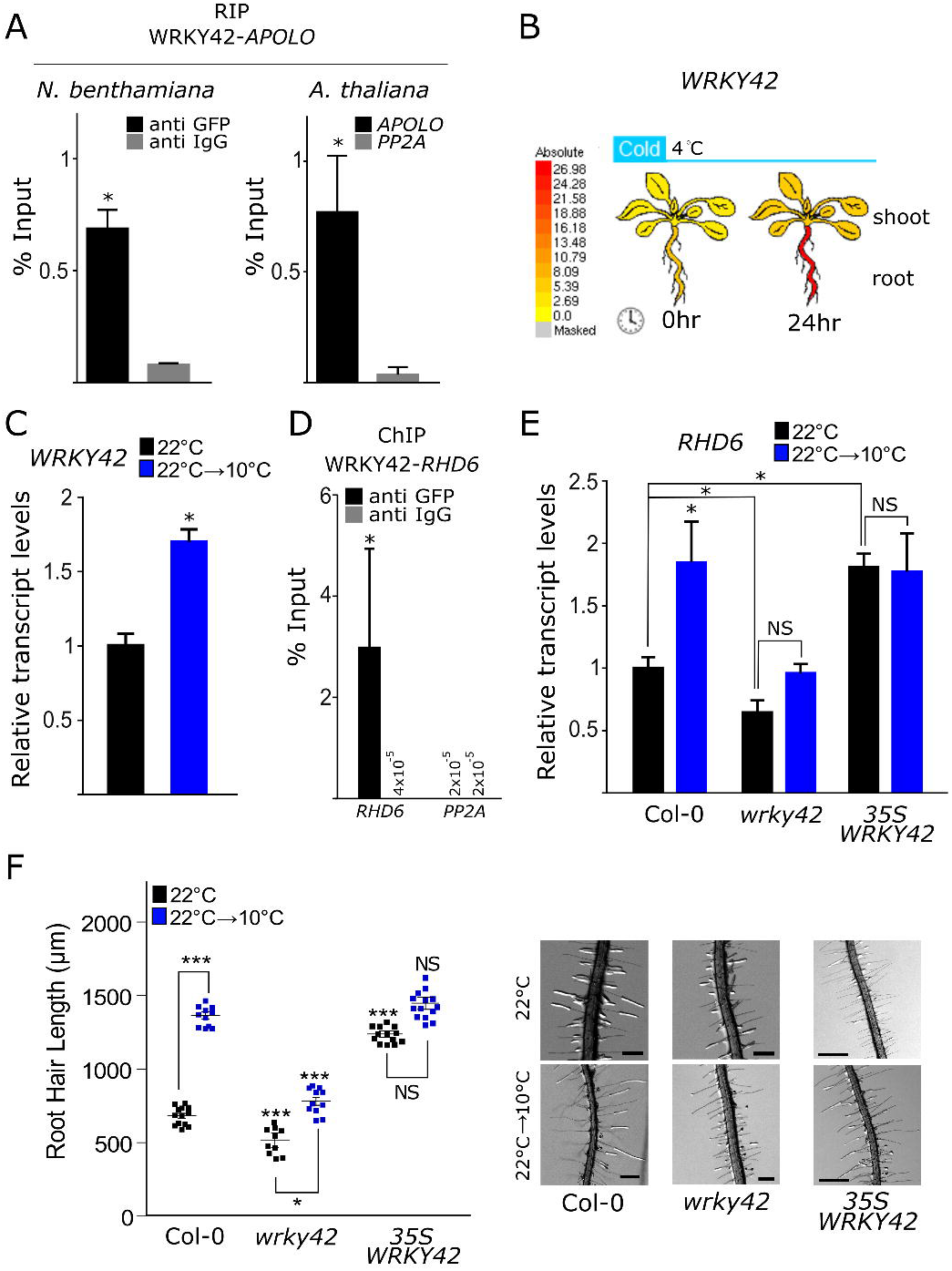
The transcription factor WRKY42 interacts with *APOLO*, directly regulates *RHD6* and participates in the response to cold. **A.** *APOLO*-WRKY42 *in vivo* interaction demonstrated by RNA Immunoprecipitation (RIP)-qPCR in transiently transformed tobacco leaves and stably transformed Arabidopsis plants. In tobacco, WRKY42:GFP and *APOLO* were co-transformed under the control of the 35S constitutive promoter. In Arabidopsis, *35S:WRKY42:GFP* transformed plants were used to detect the interaction with endogenous *APOLO*. Anti-IgG antibodies were used as a negative control. The RNA *PP2A* was considered as a RIP negative control in Arabidopsis plants. **B.** Arabidopsis eFP Browser (Waese et al., 2017) plot representing the increase of *WRKY42* transcript levels in roots of seedlings treated for 24h at 4°C. **C.** *WRKY42* transcript levels measured by RT-qPCR in roots of plants grown at 22°C treated or not for 24h at 10°C. **D.** Chromatin Immunoprecipitation (ChIP)-qPCR assay revealing regulation by WRKY42 of *RHD6* by direct recognition of its promoter region. Probes amplifying *PP2A* were used as a negative control of the experiment. Anti-IgG antibodies were used as a negative control for each pair of probes. **E.** *RHD6* transcript levels measured by RT-qPCR in roots of Col-0 plants vs. *wrky42* mutants and *35S:WRKY42:GFP* lines grown at 22°C treated or not for 24h at 10°C. **F.** Quantification of RH length of Col-0, *wrky42* and *35S:WRKY42:GFP* plants at 22°C and 10°C. Each point is the mean of the length of the 10 longest RHs identified in a single root. Representative images of each genotype are shown on the right. The scale bas represent 500μm. Asterisks indicate significant differences between Col-0 and the corresponding genotype at the same temperature (one way ANOVA followed by a Tukey-Kramer test; “*”<0.05, “***”<0.001). In A, C, D and E, the error bars represent the SD of 3 biological replicates. The asterisks indicate that the differences are significant (t test p<0.05). NS stands for no statistically significant difference. In C and E, values are normalized using the constant housekeeping transcript *PP2A*.

We thus wondered to what extent WRKY42 regulates the epigenetic landscape of the *RHD6* locus. We first observed that H3K27me3 deposition over *RHD6* is significantly augmented in the *wrky42* mutant background, in contrast to *AZG2*, an *APOLO* target non-related to WRKY42 (**Figure 4A**), consistent with reduced *RHD6* basal levels reported in *wrky42* (**Figure 3E**). Therefore, we evaluated the mutual contribution of *APOLO* and WRKY42 to their respective recognition of the *RHD6* locus. *APOLO* ChIRP-qPCR in the WT and *wrky42* mutant (**Figure 4B**) revealed similar binding to *RHD6*, indicating that WRKY42 does not participate in *APOLO*-target recognition. Reciprocally, we assessed the control of *APOLO* over WRKY42 recognition of the *RHD6* locus. To this end, we transformed transiently *A. thaliana* leaves of WT, *35S:APOLO* and RNAi-*APOLO* plants with the construct *35S:WRKY42:GFP*. Observation using confocal microscopy indicated that WRKY42:GFP is localized in the nucleus in the three genetic backgrounds (**Supplementary Figure 5**). Remarkably, WRKY42:GFP ChIP-qPCR revealed that WRKY42 binding to *RHD6* promoter is impaired both in *35S:APOLO* and RNAi-*APOLO* plants compared to WT (**Figure 4C**), hinting at a stoichiometric role of *APOLO* in the modulation of TF-chromatin interaction. To further confirm our observations, chromatin was extracted from *35S:WRKY42:GFP* seedlings and increasing amounts of *in vitro* transcribed *APOLO* were added before cross-link and regular WRKY42 ChIP over *RHD6*. Strikingly, increasing concentrations of *APOLO* gradually decoy WRKY42 away from the *RHD6* locus (**Figure 4D**), further supporting the hypothesis of a stoichiometric regulation of *APOLO* over the activity of its partner TF.

**Figure 4.**
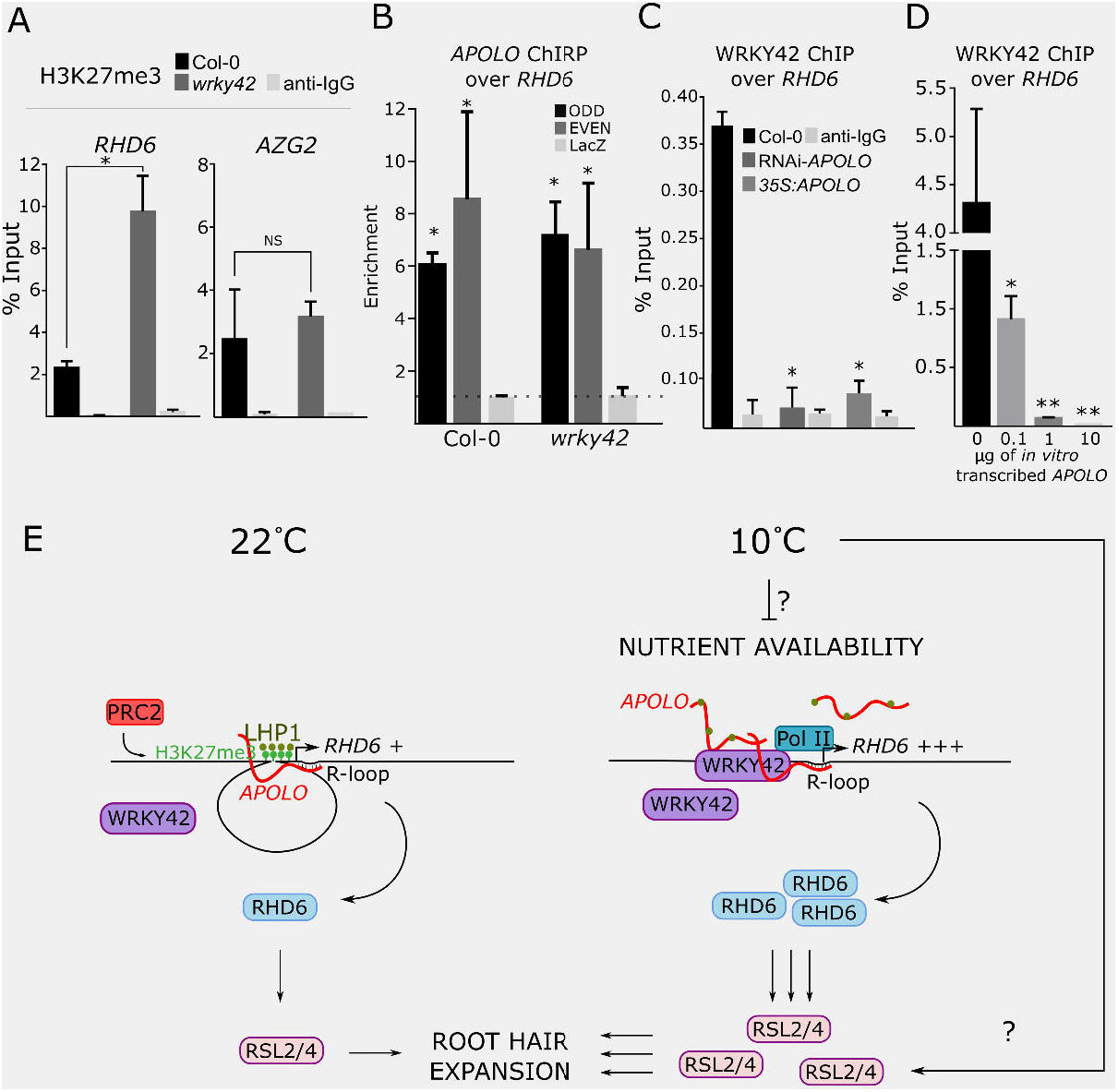
The WRKY42-*APOLO* hub activates *RHD6* transcription in response to low temperatures, promoting root hair expansion. **A.** H3K27me3 deposition in Col-0 vs. *wrky42* mutant 10-day-old seedlings grown at 23°C over *RHD6* and a WRKY42-independent *APOLO* target (*AZG2*; (Ariel et al., 2020)). An immunoprecipitation (ChIP-qPCR) was performed using an anti-IgG antibody as a negative control. **B.** *APOLO* association to DNA of the *RHD6* locus by ChIRP-qPCR in Col-0 and *wrky42* mutant plants. The background level was determined using a set of probes against LacZ RNA. **C.** WRKY42 binding to *RHD6* promoter in transiently transformed leaves of Col-0, RNAi-*APOLO* and *35S:APOLO* plants. WRKY42:GFP was immunoprecipitated using the anti-GFP antibody (ChIP-qPCR). An immunoprecipitation was performed using an anti-IgG antibody as a negative control. **D.** WRKY42 immunoprecipitation over *RHD6* with the addition of increasing concentrations of *in vitro* transcribed *APOLO* prior to cross-linking. **E.** Proposed model for the action of *APOLO* in RH growth under low temperatures. At 22 °C, *RHD6* moderate transcription is regulated by *APOLO*, H3K27me3 and LHP1, maintaining a repressive chromatin loop encompassing the *RHD6* promoter region. In response to cold, *APOLO* levels increase, still recognizing the *RHD6* locus by sequence complementarity (R-loops). Then, LHP1 (brown balls) is decoyed away from chromatin and PRC2-dependent H3K27me3 (green balls) deposition decreases in a process involving WRKY42 and leading to the opening of the *RHD6* promoter region. Additionally, *APOLO* and WRKY42 jointly activate *RHD6* transcription. RHD6 activates the transcription of downstream TF genes *RSL2* and *4* promoting RH cell expansion. Yet unknown factors activate *RSL2* and *4* independently from *APOLO* and RHD6 in response to cold. Furthermore, *APOLO* directly regulates a subset of downstream genes, likely including a subset of them controlled also by WRKY42. In A to D the error bar represents the SD of 3 biological replicates. The asterisk indicates that the difference is significant (t test “*”<0.05, “**”<0.01).

## DISCUSSION

Our results indicate that the regulation of *RHD6* expression in response to cold depends on Polycomb-dependent H3K27me3 dynamic deposition. The WRKY42-*APOLO* complex modulates the epigenetic environment of *RHD6*, activating its transcription and promoting RH growth at low temperatures. *RHD6* activation further triggers the expression of *RSL2* and *RSL4* that control the transcriptional RH program inducing cell expansion in response to cold (**Figure 4E**).

Cell fate determination in the epidermis has been extensively studied (Ryu et al., 2005; Song et al., 2011; Schiefelbein et al., 2014; Balcerowicz et al., 2015). Once trichoblast cells differentiate in the root epidermis, RHs develop as fast polar growing protuberances in response to endogenous and environmental signals (Menand et al., 2007; Yi et al., 2010; Pires et al., 2013; Marzol et al., 2017). RHs are one of the main entry points in the roots for water-soluble macronutrients, such as Pi and nitrates. Pi is an essential element for plant growth and development, and the availability of this macronutrient is a factor limiting plant productivity. In Arabidopsis roots, low Pi in the soil triggers auxin synthesis and transport, enhancing RH elongation to promote Pi uptake (Bhosale et al., 2018). Thus, auxin mediates low Pi-induced promotion of RH cell expansion. Under low soil Pi, auxin synthesis is enhanced specifically in the root cap (Stepanova et al., 2008) and transported (mostly by AUX1, PIN2, and PGP4) from the apex to the differentiation zone, specifically leading to an increase of auxin levels in trichoblasts (Jones et al., 2009; Bhosale et al., 2018; Wang et al., 2020). In response to the high-auxin microenvironment, RHs protrude from the root epidermis controlled by RHD6 and RSL1(Menand et al., 2007; Pires et al., 2013). High levels of auxin in trichoblasts trigger a signaling cascade mediated by TIR1-ARF19 (and possibly also ARF7) which directly induces the expression of *RSL4* (and likely of *RSL2*) and promote RH elongation (Yi et al., 2010; Mangano et al., 2017; Mangano et al., 2018b; Bhosale et al., 2018). ARF7 and ARF19 also activate other RH genes independently of RSL4 (Schoenaers et al., 2018). Interestingly, our results indicate that the lncRNA *APOLO* participates in the response to low temperatures. *APOLO* is directly activated by ARF7 and regulates the transcriptional activity of its neighboring gene *PINOID* (*PID*) by shaping local 3D chromatin conformation (Ariel et al., 2014; Ariel et al., 2020). *PID* encodes a kinase responsible for accurate auxin polar transport by localizing PIN2 in the root cell membrane (Friml et al., 2004). More recently, it was shown that *APOLO* can recognize a subset of distant genes across the Arabidopsis genome, most of them being related to auxin synthesis and signalling (Ariel et al., 2020). In this work, we demonstrate that a group of RH related genes is directly regulated by *APOLO* in response to cold, including the RH master regulator *RHD6*. Interestingly, *RSL2* and *RSL4* are still activated by cold in RNAi-*APOLO* roots in contrast to *RHD6* (**Figure 1C** and **Supplementary Figure 2A**), suggesting that additional yet uncovered factors, which might include ARF TFs, may trigger RH expansion in an *APOLO*/*RHD6*-independent manner, and hinting at RSL4 as a key regulator. Furthermore, we found that in addition to *RHD6*, *APOLO* directly regulates 16 RH-related genes downstream *RHD6* and *RSLs* (**Supplementary Table 1**), hinting at an intricate regulatory network controlling RH expansion. Collectively, our results uncover a lncRNA-mediated epigenetic link between environmental signals and auxin homeostasis modulating RH growth. Moreover, our observations suggest that low temperatures restrict nutrient mobility and availability in the culture medium, leading to the promotion of polar RH growth. Further research will be needed to determine what is the limiting nutrient mediating the effect of cold on RH growth.

Although substantial progress has been achieved in the identification of the molecular actors controlling RH development, the impact of chromatin conformation in the transcriptional regulation of central TFs remains poorly understood. In this study, we have revealed a new mechanism of gene regulation in RHs by which the lncRNA *APOLO* integrates chromatin-associated ribonucleoprotein complexes together with the TF WYRK42, participating in the transcriptional activation of *RHD6* and the down-stream RH gene network (**Figure 4E**). *APOLO* directly regulates the chromatin 3D conformation of the genomic region encompassing the *RHD6* locus and stoichiometrically recruits WYRK42, previously linked to Pi-starvation (Chen et al., 2009; Su et al., 2015). Low levels of *APOLO* fail to recruit WRKY42 to *RHD6* promoter region, whereas high levels of *APOLO* likely decoy WRKY42 from target chromatin, as it was shown for *APOLO* regulation over LHP1 binding activity (Ariel et al., 2020). Notably, low and high levels of *APOLO* result in higher *RHD6* basal transcriptional accumulation (through LHP1 activity and change in chromatin status), whereas both extremes impair *RHD6* activation by cold (through WRKY42 binding modulation; a model is shown in **Supplementary Figure 6**). Our results suggest that an WRKY42-*APOLO* hub regulates RH cell elongation through the master regulator RHD6, although the *APOLO*-WYRKY42 hub potentially targets several additional cell wall related genes (**Supplementary Table 1**) at the end of the pathway controlled by RHD6 and the RHD6-downstream TFs RSL2/RSL4 (Mangano et al., 2017; Mangano et al., 2018a).

Participation of epigenetic factors in root cell identity determination strongly suggests the default pattern for epidermal cell fate that can be overridden by environmental stimuli (Guimil and Dunand, 2006). Interestingly, it was reported that the expression of *GLABRA2* (*GL2*), a gene encoding a TF repressing *RHD6* in atrichoblasts, is tightly regulated at the epigenetic level. By using 3D fluorescence *in situ* hybridization, it was shown that alternative states of chromatin organization around the *GL2* locus are required to control position-dependent cell-type specification in the root epidermis (Costa and Shaw, 2006). Furthermore, *GL2* epigenetic regulation was proposed to be responsive to salt stress (Beyrne et al., 2019). In addition, a comprehensive characterization of alternative mutant lines uncovered the role of PRC2 in the regulation of RH development (Ikeuchi et al., 2015). Loss-of-function mutants in different PRC2 subunits develop unicellular RHs but fail to retain the differentiated state, generating a disorganized cell mass from each single RH. It was shown that the resulting RHs are able to undergo a normal endoreduplication program, increasing their nuclear ploidy, although they subsequently reinitiate mitotic division and successive DNA replication. It was proposed that aberrant RH development in PRC2 related mutants is due to the epigenetic deregulation of key regulatory genes such as *WOUND INDUCED DEDIFFERENTIATION 3* (*WIND3*) and *LEAFY COTYLEDON 2* (*LEC2*) (Ikeuchi et al., 2015). Here, we showed that the single mutants *clf* (PRC2) and *lhp1* (PRC1) are affected in RH growth. In addition, we found that H3K27me3 deposition and LHP1 binding to the *RHD6* locus is modulated by cold. Moreover, we showed that in the *clf* background, H3K27me3 deposition throughout the *RHD6* locus is partially impaired, and that the *clf* mutant is affected in RH elongation promoted by cold. Notably, basal transcriptional levels of *RHD6* are higher in the *clf* mutant (**Supplementary Figure 4C**). A phenotypic characterization revealed that RH density is not altered in *clf* or *lhp1* mutants, nor in *APOLO* and *WRK42* deregulated lines (**Supplementary Figure 7**), suggesting that *RHD6* is not ectopically expressed in epidermic cells. Thus, *RHD6* over-accumulation may occur in inner cell layers of the root or in RHs, although their elongation in the *clf* mutant may be blocked by additional perturbations of Polycomb-associated regulation of downstream genes. Altogether, our results suggest that Polycomb proteins participate in the control of RH-related genes transcriptional reprogramming at low temperatures.

Notably, CLF and LHP1 were shown to interact with a subset of lncRNAs in *Arabidopsis*, modulating the activity of PRC target genes (Lucero et al., 2020). Interestingly, several lncRNAs have been linked to the control of transcription in response to cold. *FLOWERING LOCUS C* is regulated by at least three lncRNAs. First, the alternative splicing of a set of antisense transcripts, collectively named as *COOLAIR*, depends on the prolonged exposure to cold, epigenetically repressing *FLC* (Marquardt et al., 2014). The use of the *COOLAIR* proximal poly(A) site results in down-regulation of *FLC* expression in a process involving FLOWERING LOCUS D (FLD), an H3K4me2 demethylase (Marquardt et al., 2014). A second lncRNA called *COLD ASSISTED INTRONIC NONCODING RNA* (*COLDAIR*) is fully encoded in the sense strand of the first intron of *FLC.* Similar to *COOLAIR*, its accumulation oscillates in response to low temperatures. It was proposed that *COLDAIR* recruits the PRC2 component CLF to target *FLC* for H3K27me3 deposition (Heo and Sung, 2011). More recently, a third lncRNA modulating *FLC* transcription was identified (Kim and Sung, 2017). The cold-responsive lncRNA *COLDWRAP* is derived from the *FLC* proximal promoter and it also interacts with PRC2. It was suggested that *COLDWRAP* functions in cooperation with the lncRNA *COLDAIR* to retain Polycomb at the *FLC* promoter through the formation of a repressive intragenic chromatin loop (Kim and Sung, 2017). Another lncRNA named *SVALKA* was shown to mediate the response to low temperatures (Kindgren et al., 2018). Interestingly, the activation of *SVALKA* by cold triggers the transcription of a cryptic downstream lncRNA, which overlaps the antisense locus of the *C-repeat/dehydration-responsive element Binding Factor 1* (*CBF1*), involved in the early response to cold in Arabidopsis. Antisense transcription causes Pol II head-to-head collision modulating transcriptional termination of *CBF1* (Kindgren et al., 2018). Here, we show that the auxin-responsive lncRNA *APOLO* is also transcriptionally modulated by cold. The differential accumulation of 24nt siRNAs across the *APOLO* locus at low temperatures indicates that this activation is related to a decrease in RdDM. Moreover, we showed here that the intergenic region between *PID* and *APOLO* acting as a divergent promoter is also activated at low temperatures in RHs, as revealed by the reporter gene *GFP*. Thus, the lncRNA *APOLO* integrates external signals into auxin-dependent developmental outputs in Arabidopsis.

In the last decade, lncRNAs have emerged as regulators of gene expression at different levels, ranging from epigenetics to protein modifications and stability (Ariel et al., 2015). Notably, it has been shown in animals that noncoding transcripts can be recognized by TFs. In humans, it was proposed that the interaction with the lncRNA *SMALL NUCLEOLAR RNA HOST GENE 15* (*SNHG15*) stabilizes the TF Slug in colon cancer cells. It was shown that *SNHG15* is recognized by the zinc finger domain of Slug preventing its ubiquitination and degradation in living cells (Jiang et al., 2018). Also, the transcriptional activity of the human gene *DIHYDROFOLATE REDUCTASE* (*DHFR*) is regulated by a lncRNA encoded in its proximal promoter. It was proposed that the nascent noncoding transcript forms a hybrid with its parent DNA and decoys the regulatory TF IIB away from the *DHFR* promoter, dissociating the transcriptional pre-initiation complex in quiescent cells (Martianov et al., 2007). The lncRNA *P21 ASSOCIATED ncRNA DNA DAMAGE ACTIVATED* (*PANDA*) was identified in human cancer and it was activated in response to DNA damage (Hung et al., 2011). *PANDA* is transcribed from the promoter region of the *CDKN1A* gene and interacts with the TF NF-YA to limit the expression of pro-apoptotic genes. The activity of *PANDA* has been linked to the progression of different tumors (Kotake et al., 2016; Shi et al., 2019). Interestingly, it was shown that in addition to NF-YA, *PANDA* interacts with the scaffold-attachment-factor A (SAFA) as well as PRC1 and PRC2 to modulate cell senescence. In proliferating cells, SAFA and *PANDA* recruit Polycomb components to repress the transcription of senescence-promoting genes. Conversely, the loss of SAFA–PANDA– PRC interactions allows expression of the senescence program (Puvvula et al., 2014). In this work, we showed that the PRC1-interacting lncRNA *APOLO* can also be recognized by the TF WRKY42, hinting at general lncRNA-mediated mechanisms linking Polycomb complexes with the transcriptional machinery across kingdoms. Furthermore, our observations indicate that the deregulation of *WRKY42* affects the epigenetic environment of *RHD6*. It was previously shown that the addition of *in vitro* transcribed *APOLO* to RNAi-*APOLO* chromatin extracts was able to partially restore R-loop formation over *APOLO* target genes, and that high levels of *APOLO* may titer LHP1 away from chromatin (Ariel et al., 2020). Here we show that the relative accumulation of the lncRNA *APOLO* can modulate the binding activity of its partner TF to common target genes. Collectively, our results strongly support that environmentally controlled cell fate in *Arabidopsis* relies on a transcriptional reprogramming governed by a network of epigenetic regulatory complexes, lncRNAs, TFs and effector proteins.

## MATERIALS AND METHODS

### Plant Material and Growth Conditions

All the *Arabidopsis thaliana* lines used were in the Columbia-0 (Col-0) background*. WRKY42* over expression transgenic plants were generated through *Agrobacterium tumefaciens* (strain EHA105)-mediated transformation (Clough and Bent, 1998). *35S:WRKY42:GFP* transformant lines were selected on MS/2 medium supplemented with kanamycin (40μg/mL) and *WRKY42* expression levels were measured by RT-qPCR (primers used are listed in **Supplementary Table 3**). The *wrky42* mutant line belongs to the SALK collection (SALK_121674C), as the one previously characterized (Su et al., 2015).The *rhd6-3/rsl1-1/rsl4-1*; *rsl2*; *rsl4*; *rsl2rsl4*, *35S:RSL4 and EXP7p:RSL4* transgenic lines were previously described and characterized (Yi et al., 2010; Hwang et al., 2017) Homozygous plants were obtained in our laboratory and genotyped using the oligonucleotides indicated in **Supplementary Table 3**. Seeds were surface sterilized and stratified at 4°C for 2d before being grown under long day conditions (16h light, 140μE.m^−2^.sec^−1^/8h dark), on ½-strength Murashige and Skoog media (1/2 MS) (Duchefa, Netherlands) with 0.8% plant agar (Duchefa, Netherlands).

### Cloning procedure

The coding region of *WRKY42* (AT4G04450) excluding the STOP codon was amplified by PCR, cloned into the Gateway entry vector pENTR/D-TOPO (Invitrogen) and recombined by Gateway technology (LR reaction) into the pK7FWG2,0 vector containing a *p35S-GFP* cassette (http://www.psb.ugent.be/gateway/index.php).

### Root hair phenotype characterization

For quantitative analyses of RH phenotypes, 10 fully elongated RH from the root elongation zone of 15-20 roots, were measured on the same conditions for each particular case and grown on vertical plates with ½-strength Murashige and Skoog media (1/2 MS) (Duchefa, Netherlands) and 0.8% plant agar (Duchefa, Netherlands) for 5 days at 22° and 3 days at 10°C. Measurements were made after 8 days. Images were captured with an Olympus SZX7 Zoom Stereo Microscope (Olympus, Japan) equipped with a Q-Colors digital camera and software Q Capture Pro 7(Olympus, Japan). RH density was determined as the number of hairs in a representative area of the root elongation zone using the same setup stated above and ImageJ software. Results were expressed as the mean ± standard error (SE) and in the case of RH density, values were expressed per mm^2^. All measurements indicate the average of three independent experiments, each involving 15-20 seedlings.

### Confocal microscopy analysis of root hairs

Confocal laser scanning microscopy was performed using Zeiss LSM5 Pascal (Zeiss, Germany) and a 40x water-immersion objective, N/A=1.2. Fluorescence was analyzed by using 488 nm laser for GFP excitation (Laser Intensity: 70%, Detector Gain:550, Amplifier Offset:0.1, Amplifier Gain:1), and emitted fluorescence was recorded between 490 and 525nm for GFP tag. Z stacks were done with an optical slice of 1μm, and fluorescence intensity was measured in 15μm ROI (Region Of Interest) at the RH tip and summed for quantification of fluorescence using ImageJ. Five replicates for each of ten roots and 15 hairs per root were observed. Col-0 wild type root hairs were used as a negative control, to check autofluorescence signal occurrence and no signal were detected in the wavelengths range stated above.

### RNA extraction and RT-qPCR

Total RNA was extracted using TRIZol (Invitrogen) and 2 μg were subjected to DNase treatment according to the manufacturer’s protocol (Thermo Scientific). One μg of DNase-free RNA was reverse-transcribed using Maxima H Minus Reverse Transcriptase (Thermo Scientific). RT-qPCR were performed using the LightCycler 480 SYBR Green I Master Kit on a LightCycler480 apparatus (Roche) using standard protocols (40 cycles, 60°C annealing). *PP2A* (AT1G13320; primers are listed in **Supplementary Table 3**) was used as reference.

### RNA Immunoprecipitation

RNA immunoprecipitation (RIP) assays were performed on transiently transformed *N. benthamiana* leaves as described in (Sorenson and Bailey-Serres, 2015), or in 10-day-old *A. thaliana 35S:WRKY42:GFP* seedlings as described in (Bardou et al., 2014), using anti GFP (Abcam ab290) and anti-IgG (Abcam ab6702). RIP was performed using Invitrogen Protein A Dynabeads. Precipitated RNAs were prepared using TRI Reagent (Sigma-Aldrich), treated with DNase (Fermentas) and subjected to RT-qPCR (High Capacity cDNA Reverse Transcription Kit (Thermo); primers used are listed in **Supplementary Table 3**). Total RNAs were processed in parallel and considered the input sample.

### Chromatin Immunoprecipitation

Chromatin immunoprecipitation (ChIP) assays were performed on 10-day-old WT seedlings treated or not during 24h at 10 °C, using anti H3K27me3 (Diagenode pAb-195-050), anti LHP1 (Covalab pab0923-P) and anti-IgG (Abcam ab6702), as described in (Ariel et al., 2020). Cross-linked chromatin was sonicated using a water bath Bioruptor Pico (Diagenode; 30sec ON/30sec OFF pulses; 10 cycles; high intensity). ChIP was performed using Invitrogen Protein A Dynabeads. Precipitated DNA was recovered using Phenol:Chloroform:Isoamilic Acid (25:24:1; Sigma) and subjected to RT-qPCR (primers used are listed in **Supplementary Table 3**). Untreated sonicated chromatin was processed in parallel and considered the input sample. For *in vitro* competition assays, *APOLO* was transcribed using the T7 transcription kit (Promega; (Ariel et al., 2020)). After regular chromatin isolation from 10-day-old *35S:WRKY42:GFP* seedlings, the sample was split in 4 independent tubes and diluted to 1ml in Nuclei Lysis Buffer without SDS. 0 μg, 0.1 μg, 1 μg and 10 μg of *APOLO* were added to each sample respectively, and incubated in rotation at 4 °C for 3h. Then, cross-linking was performed with 1% formaldehyde for 5 min at 4 °C, followed by 5 min with a final concentration of 50 mM glycine. SDS was added to a final concentration of 0.1% prior to sonication and the subsequent steps of a regular ChIP protocol.

For ChIP in transiently transformed leaves, 3-week-old *A. thaliana* were transformed as previously described (Zhang et al., 2020) In brief, *Agrobacterium tumefaciens* strain GV3101 carrying *35S:WRKY42:GFP* construct were grown for 2 days in YEB-induced medium plates at 28 °C. Agrobacterium cells were scraped and resuspended in washing solution (10 mM MgCl2, 100 μM acetosyringone). Infiltration solution (¼MS [pH = 6.0], 1% sucrose, 100 μM acetosyringone, 0.005% [v/v, 50 μl/l] Silwet L-77) was prepared with the previous solution, adjusting the OD600=0.5. The infiltration was carried out in all leaves > 0.5cm in length of between 10 and 15 plants per genotype (WT, *35S:APOLO* and RNAi-*APOLO* lines). After infiltration, plants were kept in light for 1h and then in darkness for 24h. Finally they were transferred back to light. Images and samples were obtained 3 days after infiltration. For image acquisition, infiltrated leaves were imaged with a Leica TCS SP8 confocal laser scanning microscope with excitation at 488 nm (Intensity=8%) and detection at 495-530 nm for GFP and 610-670 (gain 650) nm for chlorophyll fluorescence. Images were captured using 10X and 20X lenses, and processed using Fiji software (Schindelin et al., 2012).

### Chromatin Isolation by RNA Purification followed by qPCR or mass spectrometry

A method adapted from the ChIRP protocol (Chu et al., 2012) was developed to allow the identification of plant DNA associated to specific lncRNAs, as described in (Ariel et al., 2014; Ariel et al., 2020). Briefly, plants were *in vivo* crosslinked and cell nuclei were purified and extracted through sonication. The resulting supernatant was hybridized against biotinylated complementary oligonucleotides that tile the lncRNA of interest and lncRNA-associated chromatin was isolated using magnetic streptavidin beads. One hundred pmol of probes against *APOLO* (ODD and EVEN set of probes (Ariel et al., 2014; Ariel et al., 2020)) and the corresponding negative set against LacZ were used for independent purification. Co-purified ribonucleoprotein complexes were eluted and used to purify RNA or DNA, which were later subjected to downstream assays for quantification as previously described (Ariel et al., 2020).

The exploratory identification of plant nuclear proteins bound to *APOLO* was performed as described in (Rigo et al., 2020). Plant samples were prepared as for ChIRP-qPCR. For protein extraction, approximately 250 g of 7-day-old Col-0 plants grown on solid half-strength MS medium was irradiated three times with UV using a CROSSLINKERCL-508 (Uvitec) at 0.400 J/ cm^2^. Protein purification samples for protein extraction were DNase-treated according to the manufacturer (Thermo Scientific). After addition of 1.8ml of TCA-acetone (5ml 6.1N TCA + 45ml acetone + 35μl β-mercaptoethanol), samples were incubated overnight at −80°C. After centrifugation at 20000rpm for 20min at 4°C, the supernatant was discarded and 1.8ml of acetone wash buffer (120ml acetone, 84μl β-mercaptoethanol) was added to the samples. Then, samples were incubated 1h at −20°C and centrifuged again at 20000 rpm for 20min at 4°C. The supernatant was discarded, and the dry pellet was used for mass spectrometry analyses.

### Chromatin Conformation Capture

Chromosome conformation capture (3C) was performed basically as previously described in (Ariel et al., 2020) starting with 2g of seedlings. Digestions were performed overnight at 37°C with 400U *Dpn*II (NEB). DNA was ligated by incubation at 16°C for 5h in 4 ml volume using 100U of T4 DNA ligase (NEB). After reverse crosslinking and Proteinase K treatment (Invitrogen), DNA was recovered by Phenol:Chloroform:Isoamilic Acid (25:24:1; Sigma) extraction and ethanol precipitation. Relative interaction frequency was calculated by qPCR (primers used are listed in **Supplementary Table 3**). A region free of *Dpn*II was used to normalize the amount of DNA.

## Acknowledgments

We thank Chang Liu for providing the Hi-C plot of Figure 2B. This work was supported by grants from ANPCyT (PICT2016-0132 and PICT2017-0066) and Instituto Milenio iBio – Iniciativa Científica Milenio, MINECON to JME; ANPCyT (PICT2016-0007 and PICT2016-0289) and Fima Leloir Award to FA; UNL (CAI+D 2016) to LL; Laboratoire d’Excellence (LABEX) Saclay Plant Sciences (SPS; ANR-10-LABX-40) to MC; and CNRS (Laboratoire International Associé NOCOSYM) to MC and FA. LL, FI, JME and FA are researchers of CONICET; MM, JMP, CFF and JRM are fellows of the same institution. NM is a fellow of ANPCyT.

## Author contributions

MM, JMP, LL, CFF, JRM, NM, AC, MB and FA performed the experiments. JB, MB, FI, MC, JE and FA analyzed the data. JE and FA conceived the project. FA, JE and CFF wrote the manuscript with the contribution of all authors.

