## Supplemental Information for "The lncRNA *APOLO* interacts with the transcription factor WRKY42 to trigger root hair cell expansion in response to cold"

Moison et al., 2021: Supplemental Information file

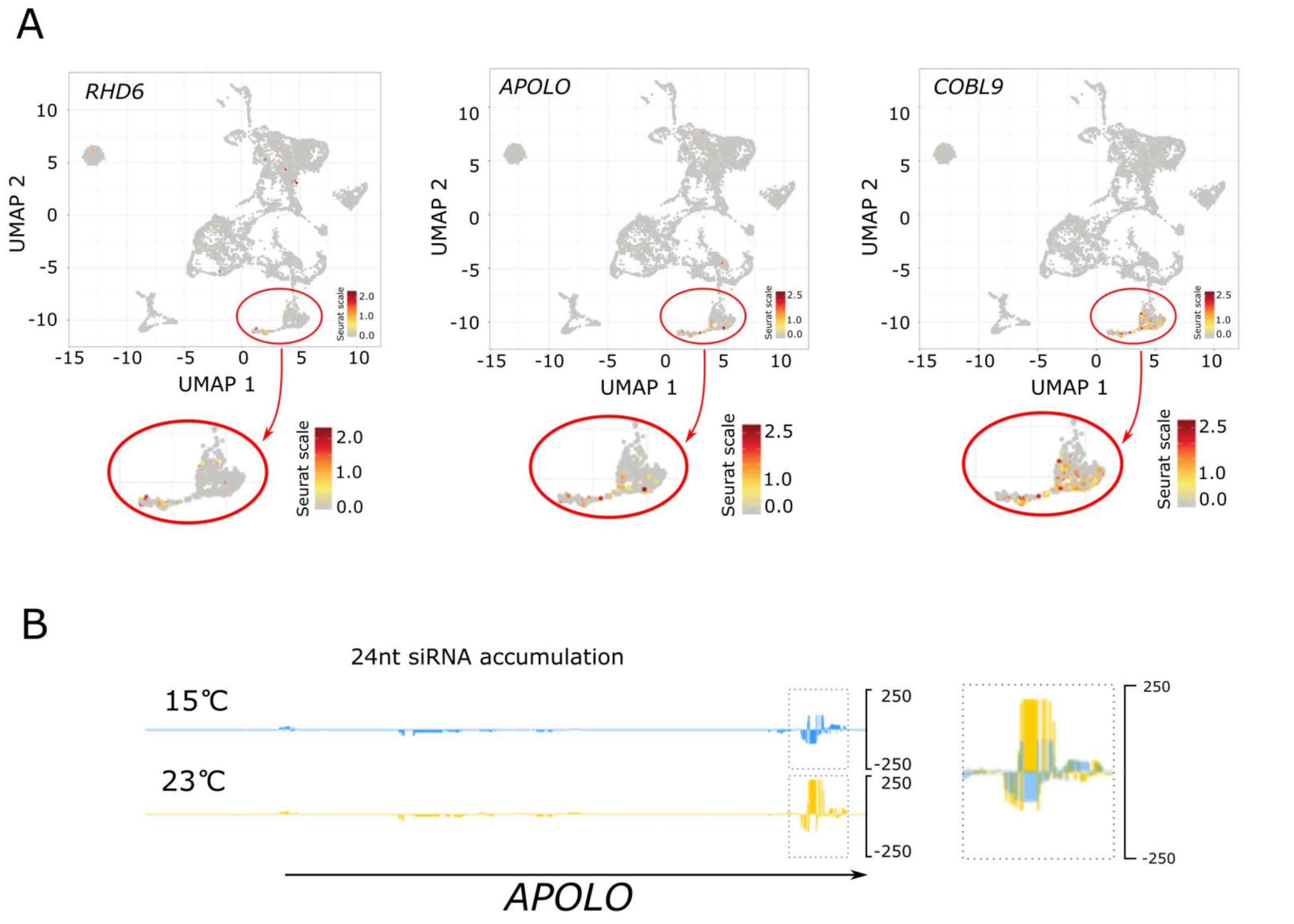


**Supplementary Figure 1: *RHD6* and *APOLO* transcripts are enriched in root hair cells**

**A.** Uniform Manifold Approximation and Projection (UMAP) clustering map of Arabidopsis root single cell RNA-seq ((Zhang et al., 2019); http://wanglab.sippe.ac.cn/rootatlas/) data. *RHD6*, *APOLO* and the trichoblast cell marker *COBL9* are respectively indicated in red in each plot.

**B.** 24nt siRNA accumulation over the *APOLO* locus revealed by small RNA-Seq of roots from Col-0 seedlings grown at 15 °C or 23 °C (Gyula et al., 2018). On the right, a zoom of the 3' region is shown, with the overlapped signal at both temperatures.

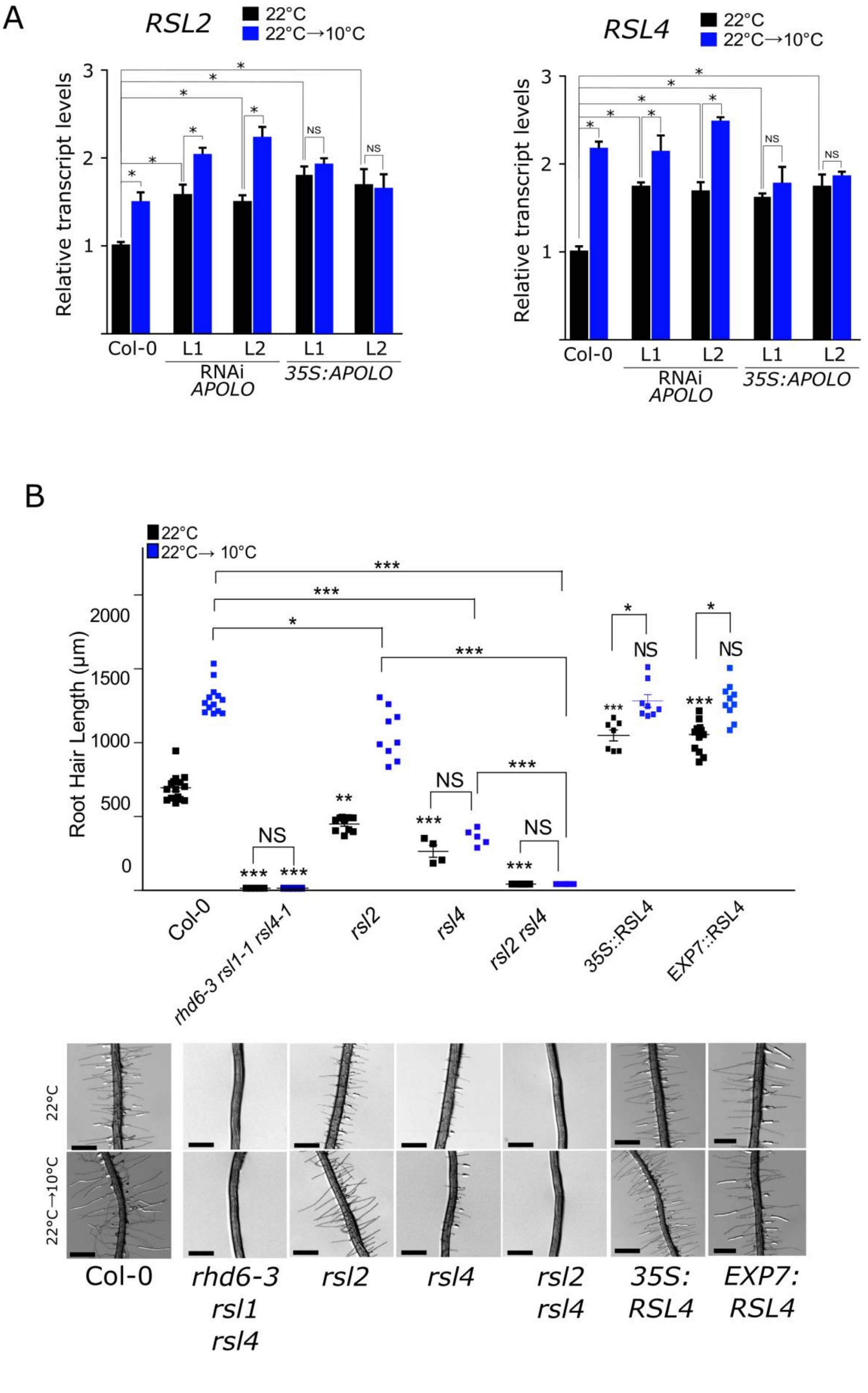


**Supplementary Figure 2. Root hair expansion in response to cold is mediated by *RSL2* and *RSL4***

**A.** *RSL2* and *RSL4* transcript levels measured by RT-qPCR in roots of Col-0 plants vs. two independent RNAi-*APOLO* lines and two independent *35S:APOLO* lines grown at 22 °C treated or not for 24h at 10 °C. The asterisks indicate that the differences are significant (t test p<0.05). NS stands for no statistically significant difference. Values are normalized using the constant housekeeping transcript *PP2A*.

**B.** Quantification of RH length of plants Col-0, *rhd6-3, rsl1/4; rsl2; rsl4; rsl2/4*; *35S:RSL4* and *pEXP7:RSL4* at 22°C and 10°C. Each point is the mean ±error of the length of the 10 longest RHs identified in the root elongation zone in a single root. Representative images of each genotype are shown below the graph. The scale bas represent 500µm. Asterisks indicate significant differences between Col-0 and the corresponding genotype at the same temperature or significant differences between temperature treatments on the same genotype at the same temperature (one way ANOVA followed by a Tukey-Kramer test; “*”<0.05, “**”<0.01, “***”<0.001), unless indicated differently. NS stands for no statistically significant difference.

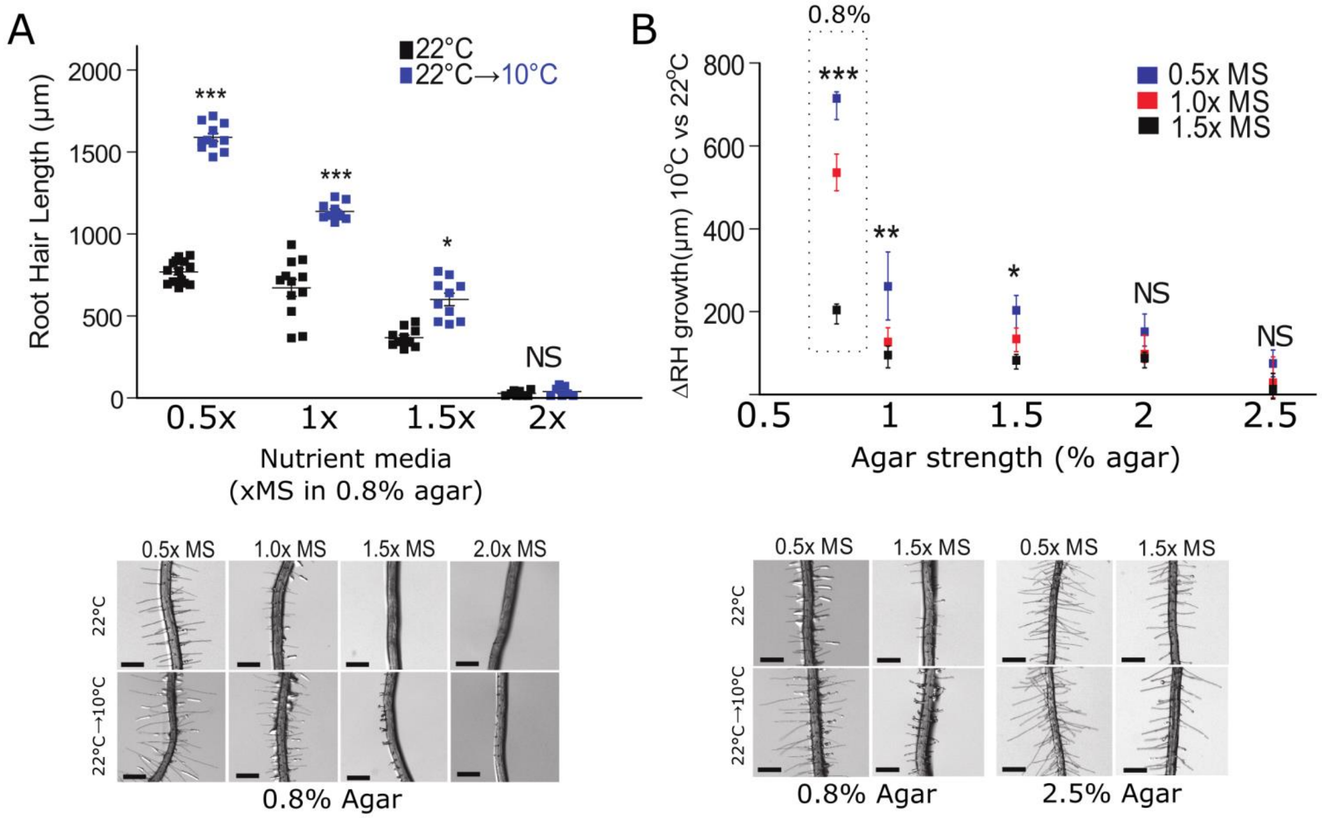


**Supplementary Figure 3. Root hair expansion in response to cold seems to be related to nutrient availability**

**A.** Quantification of RH length of Col-0 plants at 22°C and 10°C at growing concentrations of Murashige and Skoog (MS) culture media. Each point is the mean ±error of the length of the 10 longest RHs identified in the root elongation zone in a single root. Representative images of each genotype are shown below. Asterisks indicate significant differences between temperature treatments on a specific nutrient media condition (one way ANOVA followed by a Tukey-Kramer test; "*"<0.05, "***"<0.001). NS stands for no statistically significant difference.

**B.** Quantification of RH length of Col-0 plants expressed as the ratio between 10°C and 22°C at growing concentrations of MS culture media, for given concentrations of agar. Representative images of each genotype are shown below the graph. Each point is the mean ±error of the length of the 10 longest RHs identified in the root elongation zone in a single root. Note that at higher concentrations of agar, the ΔRH between conditions is lower, but the RHs are longer (as shown in the representative photos below). Asterisks indicate significant differences between agar percentage conditions (one way ANOVA followed by a Tukey-Kramer test; "*"<0.05, "**"<0.01, "***"<0.001). NS stands for no statistically significant difference. The dotted box indicates the same results as in panel A, expressed as ΔRH growth instead of absolute RH length at 22°C and 10°C independently. In the photos in A and B, the scale bas represent 500µm.

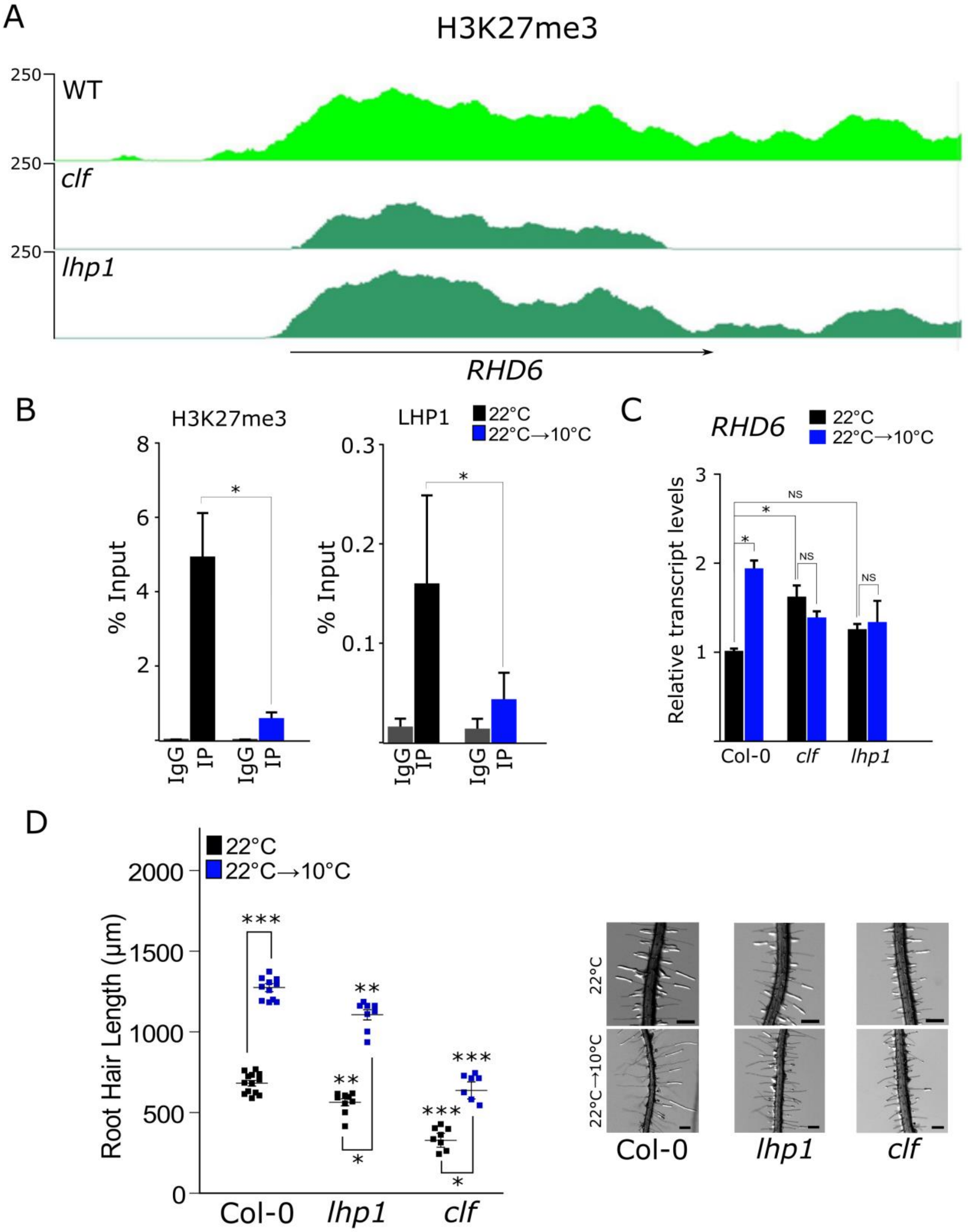


**Supplementary Figure 4: Induction of root hair elongation by low temperatures requires epigenetic Polycomb-mediated reprogramming**

**A.** H3K27me3 deposition across the *RHD6* locus in Col-0, *clf* and *lhp1* mutants, revealed by ChIP-seq from seedlings grown at 23°C (Veluchamy et al., 2016).

**B.** H3K27me3 deposition measured by ChIP-qPCR over the *RHD6* locus in Col-0 plants grown at 22°C treated or not for 24h at 10°C. The negative control was performed using an anti-IgG antibody.

**C.** *RHD6* transcript levels measured by RT-qPCR in roots of Col-0 plants vs. *clf* and *lhp1* mutants grown at 22 °C treated or not for 24h at 10 °C.

**D.** Quantification of RH length of plants Col-0, *lhp1* and *clf* at 22°C and 10°C. Each point is the mean ±error of the length of the 10 longest RHs identified in the root elongation zone in a single root. Representative images of each genotype are shown on the right. The scale bas represent 500µm. Asterisks indicate significant differences between Col-0 and the corresponding genotype at the same temperature or significant differences between temperature treatments on the same genotype at the same temperature (one way ANOVA followed by a Tukey-Kramer test; "**"<0.01, "***"<0.001).

In B and C, the error bars represent the SD of 3 biological replicates. The asterisks indicate that the difference are significant (t test p<0.05).

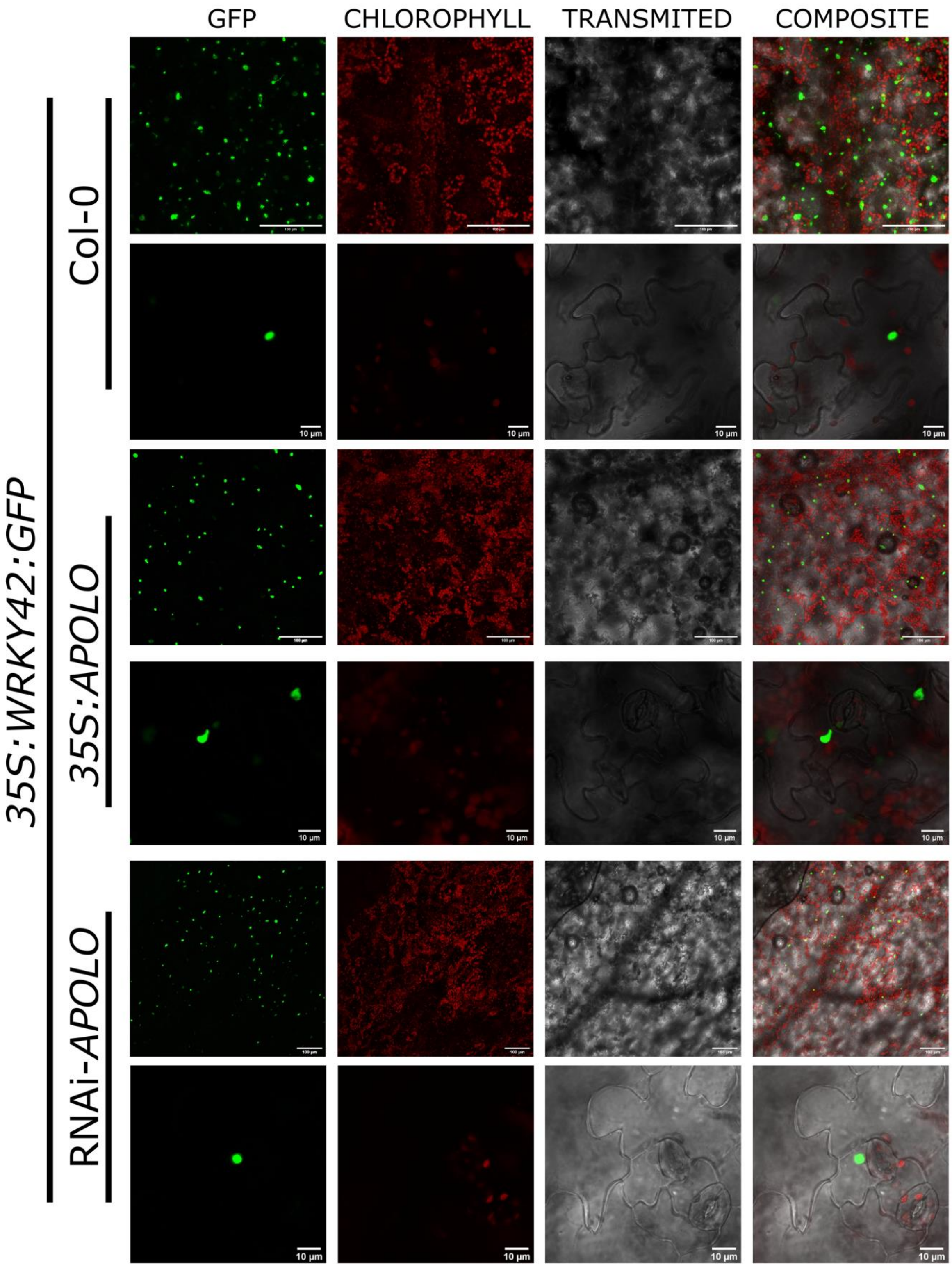


**Supplementary Figure 5: WRKY42:GFP is localized in the cell nucleus, regardless the expression levels of *APOLO***
Confocal observation of the GFP tag of WRKY42, transiently expressed in leaves of Col-0, *35S:APOLO* and RNAi-*APOLO* plants. Samples were excited at 488 nm (Intensity=8%) and detected at 495-530 nm for GFP and 610-670 (gain 650) nm for chlorophyll fluorescence. Images were captured using 10X and 20X lenses and the scale bars are indicated accordingly. The images show a high efficiency of leaf transformation for every genetic background.

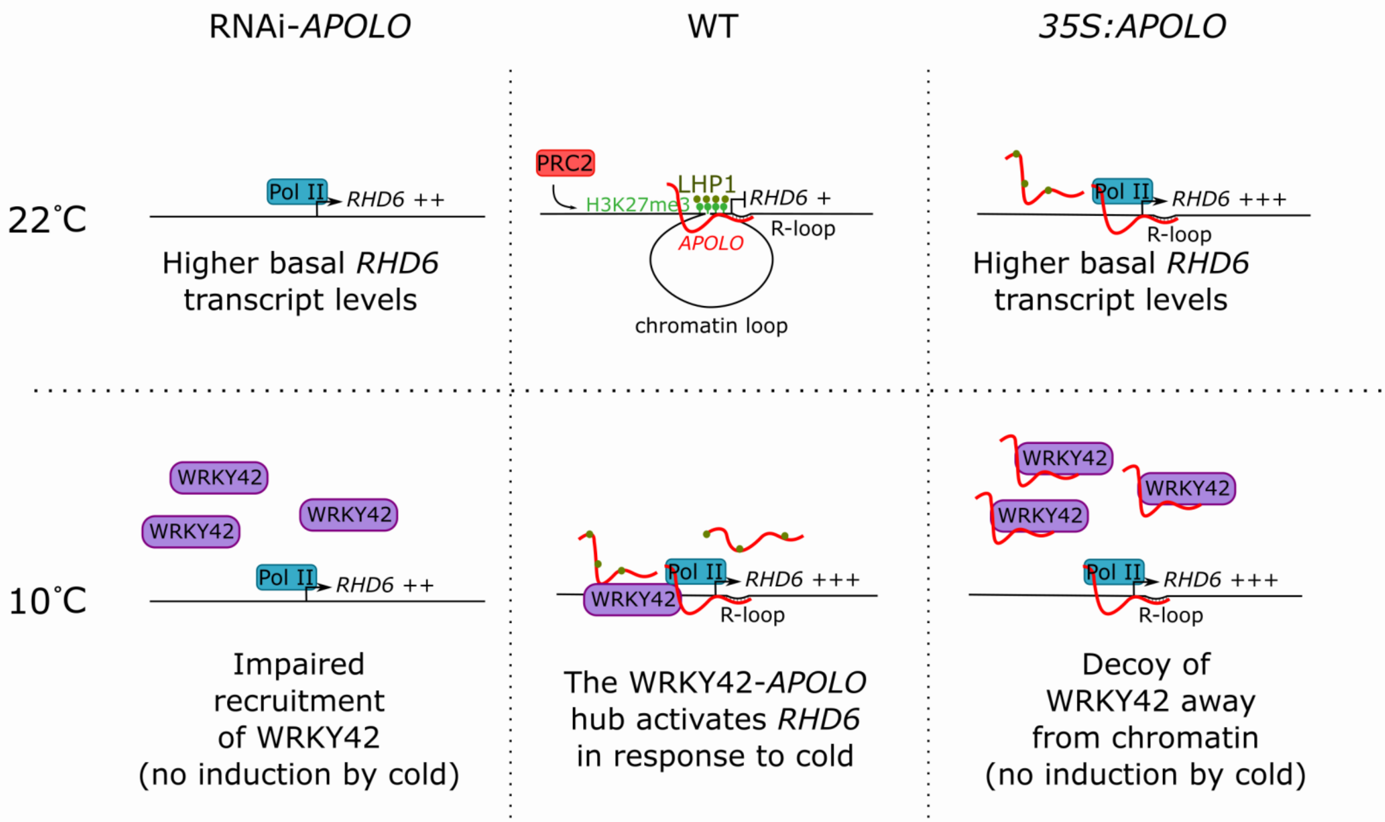


**Supplementary Figure 6: Stoichiometric role of *APOLO* revealed by the characterization of *APOLO* silencing and over-expression in transgenic plants**

Model proposed to explain the impact of *APOLO* deregulation on *RHD6* transcriptional activity. Low and high levels of *APOLO* (RNAi-*APOLO* or *35S:APOLO* lines respectively) hinder LHP1 binding and trigger the opening of the chromatin loop, enhancing *RHD6* basal expression. However, low levels or *APOLO* fail to recruit WRKY42 in response to cold, whereas high levels of *APOLO* may decoy the TF away from chromatin. As a result, both by silencing and over-expressing *APOLO*, the WRKY42-dependent activation of *RHD6* at low temperatures is impaired.

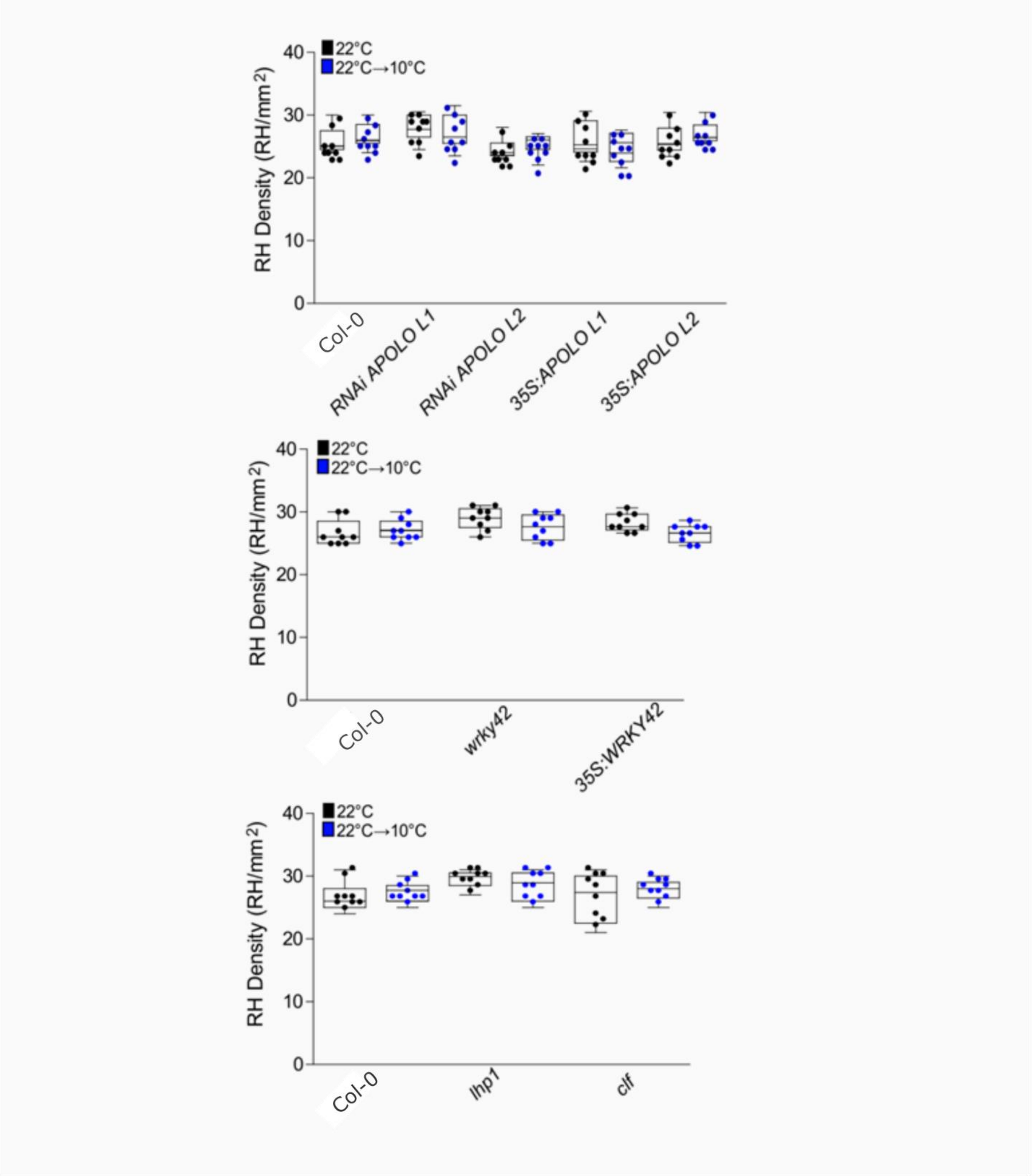


**Supplementary Figure 7: Root hair density is not altered by the deregulation of *APOLO*, *WRKY42*, *LHP1* or *CLF***
Root hair density was determined as described in the Materials and Methods section. All the lines used in this work exhibit a similar density (one way ANOVA followed by a Tukey-Kramer test; not significant) under control conditions as well as in response to cold treatment. Each point is the mean of the length of the 10 longest RHs identified in a single root.

| *APOLO* target | Gene name | LHP1 binding | H3K27me3 deposition | W-boxes in promoter | WRKY42 target (position) |
|---|---|---|---|---|---|
| **AT1G66470** | *RHD6* | YES | YES | 1 | -2521 |
| **AT1G21310** | *EXT3* | YES | YES | 4 | |
| **AT1G23720** | *EXT15* | YES | YES | 3 | |
| **AT1G26250** | *EXT18* | YES | YES | 0 | |
| **AT1G62440** | *LRX2* | YES | YES | 1 | |
| **AT2G04780** | *FLA7* | NO | NO | 2 | |
| **AT2G24980** | *EXT6* | weak | weak | 2 | |
| **AT2G26420** | *PIP5K3* | NO | NO | 2 | |
| **AT3G14310** | *PME3* | NO | NO | 0 | |
| **AT3G54580** | *EXT17* | weak | NO | 1 | |
| **AT4G08400** | *EXT7* | weak | NO | 2 | |
| **AT4G08410** | *EXT8* | weak | NO | 0 | |
| **AT4G13340** | *LRX3* | weak | NO | 3 | |
| **AT4G13390** | *EXT12* | weak | weak | 0 | |
| **AT4G40090** | *AGP3* | YES | YES | 2 | |
| **AT5G06630** | *EXT9* | YES | YES | 1 | |
| **AT5G06640** | *EXT10* | YES | YES | 0 | |

**Supplementary Table 1: *APOLO* targets related to root hair growth**
The list of *APOLO bona fide* targets was determined in (Ariel et al., 2020). The search for root hairs-related genes was performed based on (Won et al., 2009; Yi et al., 2010; Vijayakumar et al., 2016; Hwang et al., 2017). LHP1 binding and H3K27me3 deposition was investigated in previously published datasets (Veluchamy et al., 2016). W-boxes (TTGACY;(Dong et al., 2003)) present in the first 2500 bp upstream the ATG of each gene was searched manually. The position of box is given only for *RHD6*, directly recognized by WRKY42 (**Figure 3D and 4C**).

| | gene | name | locus | sample 1 | sample 2 | status | value 1 | value 2 | log2(FC) | test stat | p value | q value | significant? |
|---|---|---|---|---|---|---|---|---|---|---|---|---|---|
| *APOLO* **direct** targets (Supplementary Table 1) | AT1G66470 | ***RHD6*** | chr1:24795213-24796988 | WT | *35S:APOLO* | OK | 0,376783 | 1,05922 | 1,4912 | 2,64116 | 0,00015 | 0,00097 | yes |
| | AT1G21310 | ***EXT3*** | chr1:7453235-7455069 | WT | *35S:APOLO* | OK | 152,749 | 461,038 | 1,59372 | 6,83739 | 5,00E-05 | 0,00035 | yes |
| | AT1G23720 | ***EXT15*** | chr1:8388654-8391540 | WT | *35S:APOLO* | OK | 3,09641 | 19,245 | 2,63582 | 10,0477 | 0,00005 | 0,00035 | yes |
| | AT1G26250 | ***EXT18*** | chr1:9083837-9085464 | WT | *35S:APOLO* | OK | 3,22367 | 1,51056 | -1,09362 | -2,90259 | 0,00005 | 0,00035 | yes |
| | AT1G62440 | ***LRX2*** | chr1:23111817-23115508 | WT | *35S:APOLO* | OK | 0,292088 | 2,59795 | 3,1529 | 6,79781 | 5,00E-05 | 0,00035 | yes |
| | AT2G04780 | ***FLA7*** | chr2:1676741-1678455 | WT | *35S:APOLO* | OK | 7,18037 | 16,6557 | 1,21388 | 4,52231 | 0,00005 | 0,00035 | yes |
| | AT2G24980 | ***EXT6*** | chr2:10622452-10624132 | WT | *35S:APOLO* | OK | 0,837653 | 6,96907 | 3,05654 | 7,82993 | 5,00E-05 | 0,00035 | yes |
| | AT2G26420 | ***PIP5K3*** | chr2:11239433-11242239 | WT | *35S:APOLO* | OK | 0,077305 | 0,34491 | 2,15759 | 2,67767 | 0,0003 | 0,00181 | yes |
| | AT3G14310 | ***PME3*** | chr3:4771901-4775119 | WT | *35S:APOLO* | OK | 12,0768 | 25,3357 | 1,06894 | 4,63885 | 0,00005 | 0,00035 | yes |
| | AT3G54580 | ***EXT17*** | chr3:20200687-20203782 | WT | *35S:APOLO* | OK | 2,80301 | 22,185 | 2,98453 | 11,514 | 5,00E-05 | 0,00035 | yes |
| | AT4G08400 | ***EXT7*** | chr4:5323799-5325341 | WT | *35S:APOLO* | OK | 0,166086 | 2,09646 | 3,65795 | 5,75847 | 5,00E-05 | 0,00035 | yes |
| | AT4G08410 | ***EXT8*** | chr4:5339085-5341209 | WT | *35S:APOLO* | OK | 0,242031 | 2,59029 | 3,41985 | 6,63579 | 5,00E-05 | 0,00035 | yes |
| | AT4G13340 | ***LRX3*** | chr4:7758609-7761159 | WT | *35S:APOLO* | OK | 61,2495 | 139,492 | 1,18741 | 5,4 | 5,00E-05 | 0,00035 | yes |
| | AT4G13390 | ***EXT12*** | chr4:7783802-7785332 | WT | *35S:APOLO* | OK | 1,28902 | 9,4516 | 2,87429 | 7,90266 | 5,00E-05 | 0,00035 | yes |
| | AT4G40090 | ***AGP3*** | chr4:18580837-18581585 | WT | *35S:APOLO* | OK | 0,465013 | 6,11837 | 3,71781 | 6,1024 | 0,00005 | 0,00035 | yes |
| | AT5G06630 | ***EXT9*** | chr5:2036360-2037843 | WT | *35S:APOLO* | OK | 0,47843 | 4,28202 | 3,16191 | 6,69628 | 5,00E-05 | 0,00035 | yes |
| | AT5G06640 | ***EXT10*** | chr5:2039858-2041928 | WT | *35S:APOLO* | OK | 0,704936 | 6,53659 | 3,21297 | 8,39508 | 5,00E-05 | 0,00035 | yes |
| RH-related genes **NOT** identified as direct *APOLO* targets | AT1G02900 | ***RALF1*** | chr1:653764-654343 | WT | *35S:APOLO* | OK | 6,43265 | 11,7847 | 0,87343 | 2,3996 | 0,0001 | 0,00067 | yes |
| | AT1G12040 | ***LRX1*** | chr1:4070123-4072567 | WT | *35S:APOLO* | OK | 0,254515 | 2,57012 | 3,33601 | 6,95711 | 5,00E-05 | 0,00035 | yes |
| | AT1G12560 | ***EXPA7*** | chr1:4276518-4277899 | WT | *35S:APOLO* | OK | 0,373865 | 2,37379 | 2,6666 | 4,34024 | 5,00E-05 | 0,00035 | yes |
| | AT1G12950 | ***RSH2*** | chr1:4419769-4422638 | WT | *35S:APOLO* | OK | 0,312439 | 1,50467 | 2,2678 | 4,34933 | 5,00E-05 | 0,00035 | yes |
| | AT1G16440 | ***RSH3*** | chr1:5615814-5617672 | WT | *35S:APOLO* | OK | 0,16653 | 0,405508 | 1,28395 | 1,77649 | 0,00455 | 0,01897 | yes |
| | AT1G27740 | ***RSL4*** | chr1:9654687-9656089 | WT | *35S:APOLO* | OK | 0,105364 | 0,682308 | 2,69504 | 3,06432 | 0,0005 | 0,00284 | yes |
| | AT1G32640 | ***MYC2*** | chr1:11798809-11800988 | WT | *35S:APOLO* | OK | 22,203 | 52,6239 | 1,24496 | 5,66997 | 5,00E-05 | 0,00035 | yes |
| | AT2G35150 | ***PHI-1-like*** | chr2:14817046-14818211 | WT | *35S:APOLO* | OK | 0,264466 | 1,09189 | 2,04567 | 3,08494 | 5,00E-05 | 0,00035 | yes |
| | AT1G54970 | ***PRP1*** | chr1:20504999-20506488 | WT | *35S:APOLO* | OK | 1,07342 | 8,36667 | 2,96244 | 7,31633 | 5,00E-05 | 0,00035 | yes |
| | AT1G55330 | ***AGP21*** | chr1:20648464-20648891 | WT | *35S:APOLO* | OK | 36,0237 | 97,1117 | 1,4307 | 5,34932 | 5,00E-05 | 0,00035 | yes |
| | AT1G63450 | ***RHS8*** | chr1:23531408-23534569 | WT | *35S:APOLO* | OK | 0,03899 | 0,331918 | 3,08967 | 3,72534 | 5,00E-05 | 0,00035 | yes |
| | AT1G70460 | ***PERK13*** | chr1:26555845-26559284 | WT | *35S:APOLO* | OK | 0,258225 | 1,21635 | 2,23585 | 4,60468 | 5,00E-05 | 0,00035 | yes |
| | AT2G25260 | ***HPAT2*** | chr2:10755523-10757772 | WT | *35S:APOLO* | OK | 0,529251 | 1,33354 | 1,33324 | 2,44251 | 5,00E-05 | 0,00035 | yes |
| | AT2G43150 | ***EXT21*** | chr2:17945769-17947032 | WT | *35S:APOLO* | OK | 52,0807 | 114,204 | 1,1328 | 5,20396 | 5,00E-05 | 0,00035 | yes |
| | AT2G47540 | ***SHV2*** | chr2:19505859-19506598 | WT | *35S:APOLO* | OK | 1,0193 | 4,80441 | 2,23678 | 4,04873 | 5,00E-05 | 0,00035 | yes |
| | AT3G07340 | ***bHLH62 TF*** | chr3:2340898-2343470 | WT | *35S:APOLO* | OK | 3,61588 | 7,57866 | 1,06759 | 3,73419 | 5,00E-05 | 0,00035 | yes |
| | AT3G10710 | ***RHS12*** | chr3:3352288-3354237 | WT | *35S:APOLO* | OK | 0,11403 | 1,03108 | 3,17667 | 4,47911 | 5,00E-05 | 0,00035 | yes |
| | AT3G21180 | ***ACA9*** | chr3:7425518-7432200 | WT | *35S:APOLO* | OK | 0,058342 | 0,415952 | 2,83382 | 4,24417 | 5,00E-05 | 0,00035 | yes |
| | AT3G28550 | ***EXT16*** | chr3:10688322-10703976 | WT | *35S:APOLO* | OK | 3,33615 | 28,0079 | 3,06958 | 5,7735 | 5,00E-05 | 0,00035 | yes |
| | AT3G43960 | ***RDL3*** | chr3:15774054-15775657 | WT | *35S:APOLO* | OK | 1,63093 | 3,72675 | 1,19223 | 3,00585 | 5,00E-05 | 0,00035 | yes |
| | AT3G48340 | ***CEP2*** | chr3:17897738-17899208 | WT | *35S:APOLO* | OK | 1,69325 | 11,8683 | 2,80925 | 7,74503 | 5,00E-05 | 0,00035 | yes |
| | AT3G54580 | ***EXT17*** | chr3:20200687-20203782 | WT | *35S:APOLO* | OK | 2,80301 | 22,185 | 2,98453 | 11,514 | 5,00E-05 | 0,00035 | yes |
| | AT3G60330 | ***HA7*** | chr3:22297406-22303673 | WT | *35S:APOLO* | OK | 0,75475 | 1,83265 | 1,27986 | 3,54359 | 5,00E-05 | 0,00035 | yes |
| | AT3G62680 | ***PRP3*** | chr3:23182721-23183994 | WT | *35S:APOLO* | OK | 1,90231 | 17,1835 | 3,1752 | 9,02884 | 5,00E-05 | 0,00035 | yes |
| | AT4G02270 | ***RHS13*** | chr4:992174-993038 | WT | *35S:APOLO* | OK | 0,956108 | 7,16393 | 2,90551 | 5,80735 | 5,00E-05 | 0,00035 | yes |
| | AT4G17215 | ***Pollen Ole e 1 allergen and extensin*** | chr4:9655836-9656577 | WT | *35S:APOLO* | OK | 1,17841 | 4,76508 | 2,01566 | 3,84981 | 5,00E-05 | 0,00035 | yes |
| | AT4G22080 | ***RHS14*** | chr4:11700554-11702666 | WT | *35S:APOLO* | OK | 0,059309 | 0,757044 | 3,67406 | 3,68273 | 0,00025 | 0,00154 | yes |
| | AT4G25400 | ***bHLH TF*** | chr4:12981294-12982468 | WT | *35S:APOLO* | OK | 1,35187 | 4,32603 | 1,67809 | 3,18257 | 5,00E-05 | 0,00035 | yes |
| | AT4G29180 | ***RHS16*** | chr4:14385592-14389689 | WT | *35S:APOLO* | OK | 0,099263 | 0,521194 | 2,39249 | 3,82728 | 5,00E-05 | 0,00035 | yes |
| | AT4G29930 | ***bHLH TF*** | chr4:14644008-14647591 | WT | *35S:APOLO* | OK | 2,30654 | 7,32455 | 1,66701 | 4,31108 | 5,00E-05 | 0,00035 | yes |
| | AT4G33880 | ***RSL2*** | chr4:16239362-16241137 | WT | *35S:APOLO* | OK | 0,038579 | 0,306499 | 2,99001 | 2,43441 | 0,0083 | 0,03137 | yes |
| | AT4G36930 | ***SPT*** | chr4:17414126-17416294 | WT | *35S:APOLO* | OK | 1,36121 | 2,40537 | 0,82137 | 2,05934 | 0,00055 | 0,00309 | yes |
| | AT5G05500 | ***MOP10*** | chr5:1629666-1630396 | WT | *35S:APOLO* | OK | 0,56784 | 5,15021 | 3,18108 | 5,39453 | 5,00E-05 | 0,00035 | yes |
| | AT5G19560 | ***ROPGEF10*** | chr5:6603290-6606448 | WT | *35S:APOLO* | OK | 0,058391 | 0,406499 | 2,79944 | 2,91293 | 0,0004 | 0,00233 | yes |
| | AT5G22410 | ***RHS18*** | chr5:7426313-7427964 | WT | *35S:APOLO* | OK | 0,342718 | 3,78832 | 3,46647 | 6,35329 | 5,00E-05 | 0,00035 | yes |
| | AT5G35190 | ***EXT13*** | chr5:13434180-13435167 | WT | *35S:APOLO* | OK | 1,52769 | 15,7388 | 3,3649 | 8,58919 | 5,00E-05 | 0,00035 | yes |
| | AT5G41315 | ***GL3*** | chr5:16529454-16532879 | WT | *35S:APOLO* | OK | 0,235867 | 0,44781 | 0,92491 | 1,51906 | 0,0112 | 0,04019 | yes |
| | AT5G51060 | ***RHD2*** | chr5:20752999-20762431 | WT | *35S:APOLO* | OK | 0,213913 | 1,38566 | 2,69548 | 3,1665 | 0,0001 | 0,00067 | yes |
| | AT5G51790 | ***bHLH120 TF*** | chr5:21039510-21041446 | WT | *35S:APOLO* | OK | 2,80331 | 7,68882 | 1,45563 | 2,87659 | 5,00E-05 | 0,00035 | yes |
| | AT5G54380 | ***THE1*** | chr5:22077059-22080338 | WT | *35S:APOLO* | OK | 19,729 | 40,4626 | 1,03627 | 4,68266 | 5,00E-05 | 0,00035 | yes |
| | AT5G58010 | ***LRL3*** | chr5:23483544-23484889 | WT | *35S:APOLO* | OK | 0,163107 | 1,32569 | 3,02286 | 3,78898 | 5,00E-05 | 0,00035 | yes |
| | AT5G67400 | ***RHS19*** | chr5:26894855-26896488 | WT | *35S:APOLO* | OK | 0,319731 | 2,96103 | 3,21117 | 5,60471 | 5,00E-05 | 0,00035 | yes |

**Supplementary Table 2: Root hair-related genes deregulated in *35S:APOLO* seedlings**

The list of deregulated genes was determined by RNA-Seq (Ariel et al., 2020). The search for root hairs-related genes was performed based on (Won et al., 2009; Yi et al., 2010; Vijayakumar et al., 2016; Hwang et al., 2017). Genes are classified as directly (same as in **Supplementary Table 1**) or indirectly regulated by *APOLO*. *RHD6*, *RSL2* and *RSL4* are highlighted in red.

| | | Probes sequences | Experiment |
|---|---|---|---|
| AT1G66470 *RHD6* | F | TCCCGATTCTTCCCTTAAAAA | 3C |
| | R | AAAGCAAACCTTGGCTGCTA | |
| | F | AATGACCATCCCAACGAGAC | ChIRP, RT-qPCR |
| | R | CATAAGCTGCATCTGGCTGA | |
| | F | GGATTAGAAAGACCAGCTGCTTTAG | ChIP |
| | R | CCAGTTTGTGAACGACGACA | |
| AT2G34655 *APOLO* | F | CTTCGAGGCGCTAAACAATC | RIP, RT-qPCR |
| | R | ACAGCGGTGCCACCTATTAC | |
| AT5G50300 *AZG2* | F | GGGAGTTGCAAAATGGCTTA | ChIP |
| | R | CCGACCATGTGAAAAATTCC | |
| AT4G04450 *WRKY42* | F | CCGCTAAACGCTGGTGAGA | RT-qPCR |
| | R | CGGCAAAGTTGGGATTGG | |
| | SALK_049063_LP2 | AAGCACGCTTCGATATCTGAG | genotyping |
| | SALK_049063_RP2 | AGTTGCGACAATCAATGGATC | |
| | SALK_049063_LP4 | TTTGTGCGTCTGTTACGTACG | |
| | SALK_049063_RP4 | GTACTTGCTTGCGAACAGGAC | |
| | F | CACCATGTTTCGTTTTCCGGTAAGTC | cloning |
| | R | TTGCCTATTGTCAACGTTGCTC | |
| AT4G33880 *RSL2* | F | TCCCCAATGGAACAAAGGTC | RT-qPCR |
| | R | TCTCGGTGAGCTGAGACCAA | |
| AT1G27740 *RSL4* | F | GTGCCAAACGGGACAAAAGT | RT-qPCR |
| | R | TTGTGATGGAACCCCATGTC | |
| AT1G13320 *PP2A* | F | TAACGTGGCCAAAATGATGC | RIP, RT-qPCR |
| | R | GTTCTCCACAACCGCTTGGT | |

**ChIRP 3'BIO probes**

| ODD probes | |
|---|---|
| **number** | **sequence** |
| APOLO 1 | aaccagccaatgaacagatg |
| APOLO 3 | gacaagtcacacctacactc |
| APOLO 5 | agttccaaggaatccatacc |
| APOLO 7 | gacagcggtgccacctatta |
| APOLO 9 | ttacaacaagccactccgta |
| APOLO 11 | accaaaacaaccaaaatttt |
| APOLO 13 | ccttacaaacagagcaaagt |
| APOLO 15 | ggaagcaaagcccaaaggaa |
| APOLO 17 | caagtaaccccagaaaacta |
| APOLO 19 | gattccggtgaaatacaagg |

| EVEN probes | |
|---|---|
| **number** | **sequence** |
| APOLO 2 | gattgtttagcgcctcgaag |
| APOLO 4 | cgagaagaactaggccaaag |
| APOLO 6 | tgaagactaccttacataga |
| APOLO 8 | acaaggaacttccaaccaaa |
| APOLO 10 | caacatctcgtcaaccacat |
| APOLO 12 | cgaaactaaaacaaagaagc |
| APOLO 14 | ggaaataacaaggcaaaaca |
| APOLO 16 | ccgacgattaaaaggataat |
| APOLO 18 | gaaatacaaagccggcggtt |
| APOLO 20 | tcgtctgaaagtttattata |

| EVEN LacZ probes | |
|---|---|
| **number** | **sequence** |
| LacZ 1 | ccagtgaatccgtaatcatg |
| LacZ 5 | aatgtgagcgagtaacaacc |
| LacZ 7 | aataattcgcgtctggcctt |
| LacZ 9 | aattcagacggcaaacgact |
| LacZ 11 | atcttccagataactgccgt |
| LacZ 13 | gctgatttgtgtagtcggtt |
| LacZ 15 | aactgttacccgtaggtagt |
| LacZ 17 | tttcgacgttcagacgtagt |
| LacZ 19 | accattttcaatccgcacct |
| LacZ 21 | ttcatcagcaggatatcctg |

**Supplementary Table 3: List of oligonucleotides used in this work**
